# ConnectedReads: machine-learning optimized long-range genome analysis workflow for next-generation sequencing

**DOI:** 10.1101/776807

**Authors:** Chung-Tsai Su, Sid Weng, Yun-Lung Li, Ming-Tai Chang

## Abstract

Current human genome sequencing assays in both clinical and research settings primarily utilize short-read sequencing and apply resequencing pipelines to detect genetic variants. However, theses mapping-based data analysis pipelines remains a considerable challenge due to an incomplete reference genome, mapping errors and high sequence divergence. To overcome this challenge, we propose an efficient and effective whole-read assembly workflow with unsupervised graph mining algorithms on an Apache Spark large-scale data processing platform called ConnectedReads. By fully utilizing short-read data information, ConnectedReads is able to generate assembled contigs and then benefit downstream pipelines to provide higher-resolution SV discovery than that provided by other methods, especially in high diversity against reference and N-gap regions of reference. Furthermore, we demonstrate a cost-effective approach by leveraging ConnectedReads to investigate all spectra of genetic changes in population-scale studies.

## Background

Whole-genome sequencing (WGS) is increasingly used in biomedical research, clinical, and personalized medicine applications to identify disease- and drug-associated genetic variants in humans, all with the goal of advancing precision medicine (*1*). At present, next-generation sequencing (NGS, also called short-read sequencing (SRS)) is a well-established technology used to generate whole-genome data due to its high throughput and low cost (*2*). Resequencing, especially of human samples, is one of the popular applications of NGS. This process maps raw reads against a reference genome and determines all kinds of genomic variations, including single nucleotide polymorphisms (SNPs) and indels as well as genetic rearrangements and copy-number variants (CNVs) (*3*). However, a fundamental flaw in the resequencing pipeline is that it ignores the correlation between sequence reads; thus, resequencing does not fully and properly utilize sequence data and may generate inconsistent alignments, which make variant calling, especially structural variant (SV) calling, more complicated (*4, 5*). Since the human reference genome is incomplete and contains many low-complexity regions, assembling sequence reads without reference bias would be a proper way to overcome the above challenges. Nonetheless, assembly-based approaches for WGS data suffer from several computational challenges, such as high computing resource requirements and long turnaround times.

In this article, we propose an efficient whole-read assembly workflow with unsupervised graph mining algorithms on an Apache Spark large-scale big data processing platform called ConnectedReads. By leveraging the in-memory cluster computing framework of Apache Spark (*6*), ConnectedReads takes less than 20 hours to process 30-fold human WGS data and generates corresponding long assembled contigs for downstream analysis, such as read mapping, variant calling or phasing. The spec. of cluster and the execution time for each step of ConnectedReads are described in the method session. To evaluate the data correctness of ConnectedReads, we use 68 high-confidence insertions in the NA12878 sample detected by svclassify (*7*). To demonstrate the influence of reference incompleteness, mapping error and high sequence divergence on NGS data analysis, three public available samples from different populations are used. Through ConnectedReads, we are able to investigate unique non-reference insertions (UNIs) and non-repetitive, non-reference (NRNR) sequences from population datasets (*8, 9*). Furthermore, high resolution for SVs are provided by leveraging ConnectedReads, especially on breakpoints of insertions. In conclusion, ConnectedReads optimizes NGS reads to generate long assembled contigs, not only reducing mapping error but also benefit the downstream data analysis pipelines.

## Results

### Data preparation

To exemplify the challenges of WGS data analysis mentioned above and assess the utility of ConnectedReads, three WGS datasets from three different ethnic groups were selected from publicly available databases, as listed in Table 1, including NA12878 of European ancestry, NA24694 of Asian ancestry, and NA19240 of African ancestry. In addition, the samples were around 30X and sequenced by three different Illumina platforms, namely, the NovaSeq 6000 (NA12878), HiSeq 2500 (NA24694), and HiSeq X Ten (NA19240) platforms. Therefore, we believe that these datasets are representative of the majority of WGS data on cohort datasets and national bio-bank projects.

**Table 1.** Description of WGS datasets

| Sample | Platform | Coverage | Read Length | Description | Source |
| --- | --- | --- | --- | --- | --- |
| NA12878 | NovaSeq 6000 | 30X (313,716,127<br>read pairs) | 151 | HG001 (Population: CEU) | *a |
| NA24694 | HiSeq 2500 | 30X (311,220,694<br>read pairs) | 148 | HG006, Father of The Han<br>Chinese GIAB Trio | *b |
| NA19240 | HigSeq X Ten | 35X (376,862,646<br>read pairs) | 151 | Yoruba(Nigeria)<br>(Population: YRI) | *c |

Data sources:
\*a https://www.ebi.ac.uk/ena/data/view/ERR2438055
\*b https://www.ncbi.nlm.nih.gov/sra/SRX1388455
\*c https://www.ncbi.nlm.nih.gov/sra/SRX4637790

### Evaluations

The results for the datasets in Table 1 obtained by applying the ConnectedReads workflow with default settings are listed in Table 2. Comparing to SRS data in Table 1, ConnectedReads is clearly able to reduce the number of contigs and total base pairs by more than 96% and 87%, respectively. Take NA12878 for examples, 313, 716, 127 read pairs with length of 151 are transformed to 16,348,524 contigs (Reduction rate: 97.4%). In addition, the total BPs is reduced from 94,742,270,354 on SRS to 10,466,069,046 on ConnectedReads (Reduction Rate: 88.9%). Since ConnectedReads generates assembled contigs based on paired-end or barcode information, any two contigs would not be assembled together without sufficient supports for paired-end information or overlaps. Although ConnectedReads aims to construct more accurate assembled contigs rather than longer ones without haplotypes, it is usually able to construct several contigs of more than 30 Kbps. In addition, there are 1,402,511, 1,224,389 and 2,082,886 contigs of >=1 Kbps for NA12878, NA24694 and NA19240, respectively. Furthermore, the length of contigs is strongly correlated with coverage and read length. The deeper the coverage is, the longer the contigs that can be generated are.

**Table 2.** Description of the contigs of three datasets generated by ConnectedReads.

| Sample | NA12878 | NA24694 | NA19240 |
| --- | --- | --- | --- |
| Number of contigs | 16,348,524 | 18,900,370 | 18,261,647 |
| Total base pairs (bps) | 10,466,069,046 | 9,925,886,082 | 14,093,570,587 |
| Average length | 640 | 525 | 772 |
| Longest contig | 37,619 | 33,904 | 32,145 |
| # of singletons (<=151) | 8,351,671 | 11,305,117 | 8,355,394 |
| >=1 Kbps | 2,899,483 | 2,470,630 | 4,487,568 |
| >=2 Kbps | 1,402,551 | 1,224,389 | 2,082,886 |
| >=5 Kbps | 263,878 | 256,178 | 277,357 |

ConnectedReads offers two advantages for downstream analysis. One is mapping recovery, and the other is SV detectability. Both of these advantages are described comprehensively by the following experimental results and case studies.

First, mapping recovery is the best way to demonstrate the advantage of ConnectedReads. According to the evolution of NGS technology in recent years, longer reads can reduce two kinds of mapping errors, namely, false mapping and uncovered regions based on the reference genome (*10*). However, it is hard to determine whether a mapping record is falsely aligned because of several complicated situations, such as sequencing errors, an incomplete reference genome, high sequence divergence and SVs (*5*). Therefore, the recovery rate is a proper measurement for evaluating the performance of ConnectedReads and SRS. Mapping recovery means all of positions on human reference is not mapped by any SRS data but able to be covered by ConnectedReads data. For example, the uncovered regions of the reference genome means that there is no short reads mapped by using Genome Analysis Toolkit (GATK) Best Practices (i.e., BWA-MEM) but contigs of ConnectedReads are mapped on some of the regions by using minimap2 (*11*). These regions are called as mapping recovery. In terms of NA12878 mapping recovery, as illustrated in 0, there is a 15%-90% recovery rate for each chromosome on hg38 (excluding chrY). For example, chr1 has 319,018 uncovered bps with SRS, but 135,253 bps (42.4%) can be recovered by ConnectedReads. The best recovery rate (89.6%) is on chr21, and the worst recovery rate (15.4%) is on chr17. Large regions are often recovered when large deletions occur. Overall, ConnectedReads is able to reconstruct the mapping information for uncovered regions in the reference genome by using SRS data, and it may be the best candidate complementary to Illumina short-read data.

Second, SV detection remains a challenge in SRS (*12*). Using ConnectedReads technology will significantly mitigate this challenge because ConnectedReads has the similar advantage as long-read sequencing (LRS) from Nanopore and PacBio. The longer the read is, the more correct the mapping result is and the more easily SVs can be detected. Fig. 2 shows the numbers of insertions and deletions identified in NA12878. The statistical method is simply based on a CIGAR string generated by minimap2. When the threshold of mapping quality (MAPQ) is increased, fewer insertions and detections are identified. Since several alignment records with an MAPQ of 0 were falsely mapped in our investigation, the threshold of MAPQ was set to 1 in the following experiments to balance the precision and sensitivity of SV detection. In addition, several recent studies have revealed that every human genome has approximately 20,000 SVs that span at least 10 million bps (*13, 14*). ConnectedReads identifies 21,855 SVs, and the number of SVs is similar to that obtained in previous studies (*13, 14*).

**Fig. 1.**
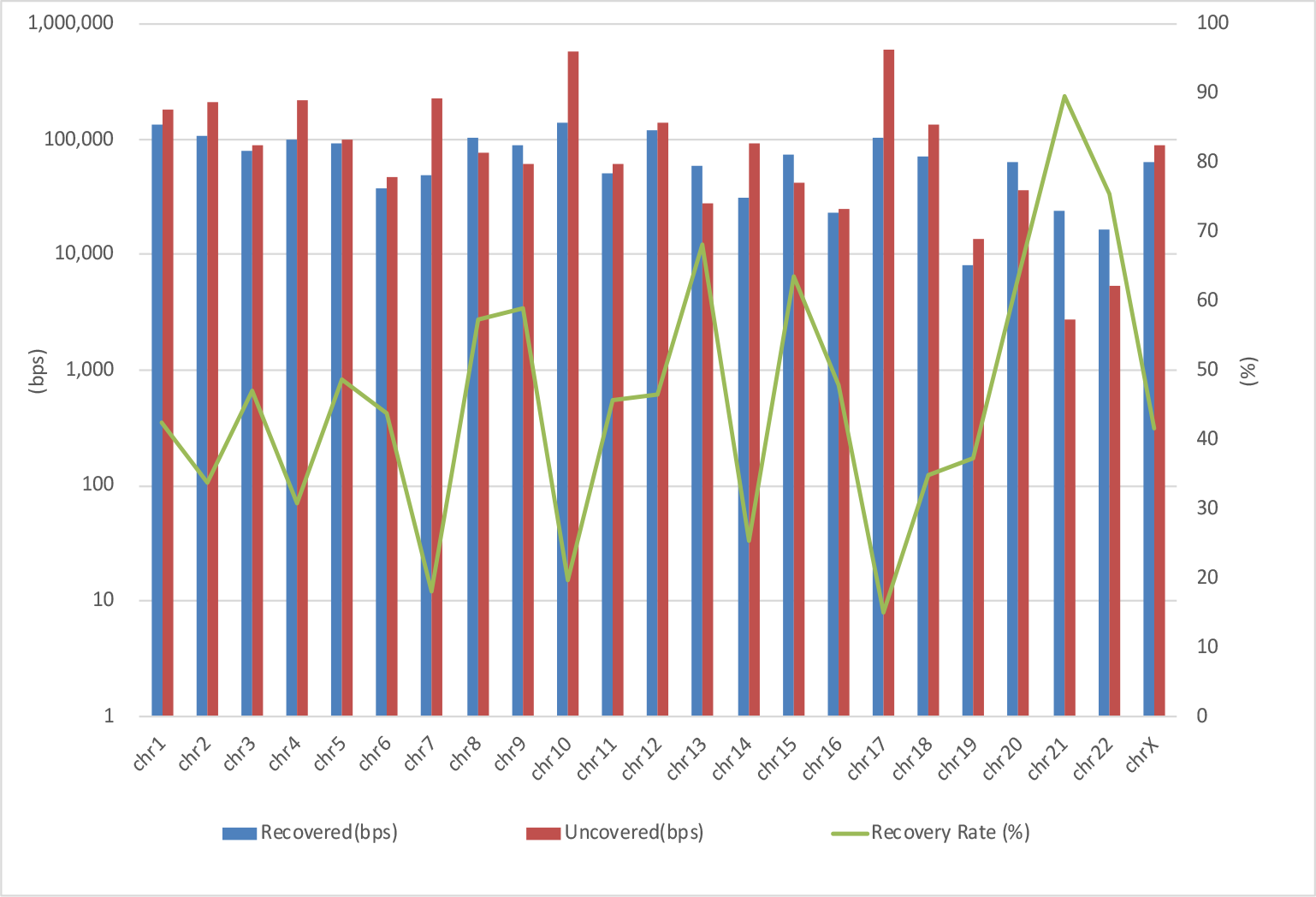
Mapping recovery of NA12878 (short reads vs. ConnectedReads). BWA and minimap2 are adopted for the short-read data and ConnectedReads contigs, respectively. Recovery Rate = # recovered / (# recovered + # uncovered).

**Fig. 2.**
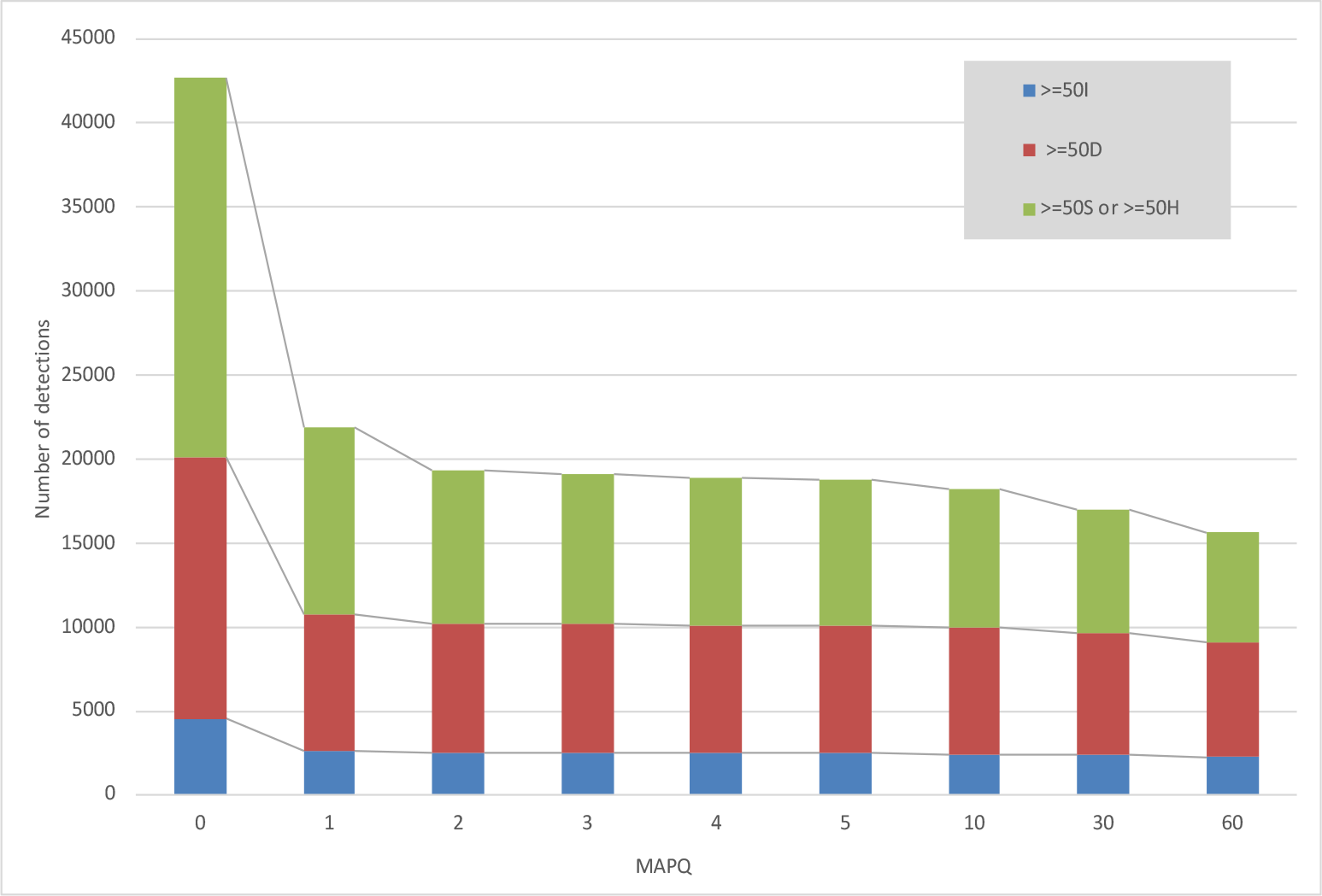
Number of SV vs. MAPQ (NA12878). The priority order used for counting is insertion > deletion > soft-/hard-clipped sequence. Any two SVs will be merged if their distance is less than 50 bps. This means that if one insertion and one deletion are encountered in the same location, only one insertion is counted.

Furthermore, an interesting phenomenon on Fig. 3 is observed when the insertions and deletions shown in Fig. 2 are sorted by length. Based on the 1000 Genomes Project and several studies (*15, 16*), the number of SVs decreases as the length of the SVs increases. Therefore, the majority of the SVs are small indels (<50 bps). Then, the trend of the distribution slightly decreases as the length of the SVs increases, except for the peak at 250-300 bps. The reason is due to abundant Alu elements whose body lengths are approximately 280 bases (*17*). In addition, several studies have reported this phenomenon when using PacBio LRS (*13*). Therefore, ConnectedReads is able to complement SRS in not only mapping recovery but also SV detection.

**Fig. 3.**
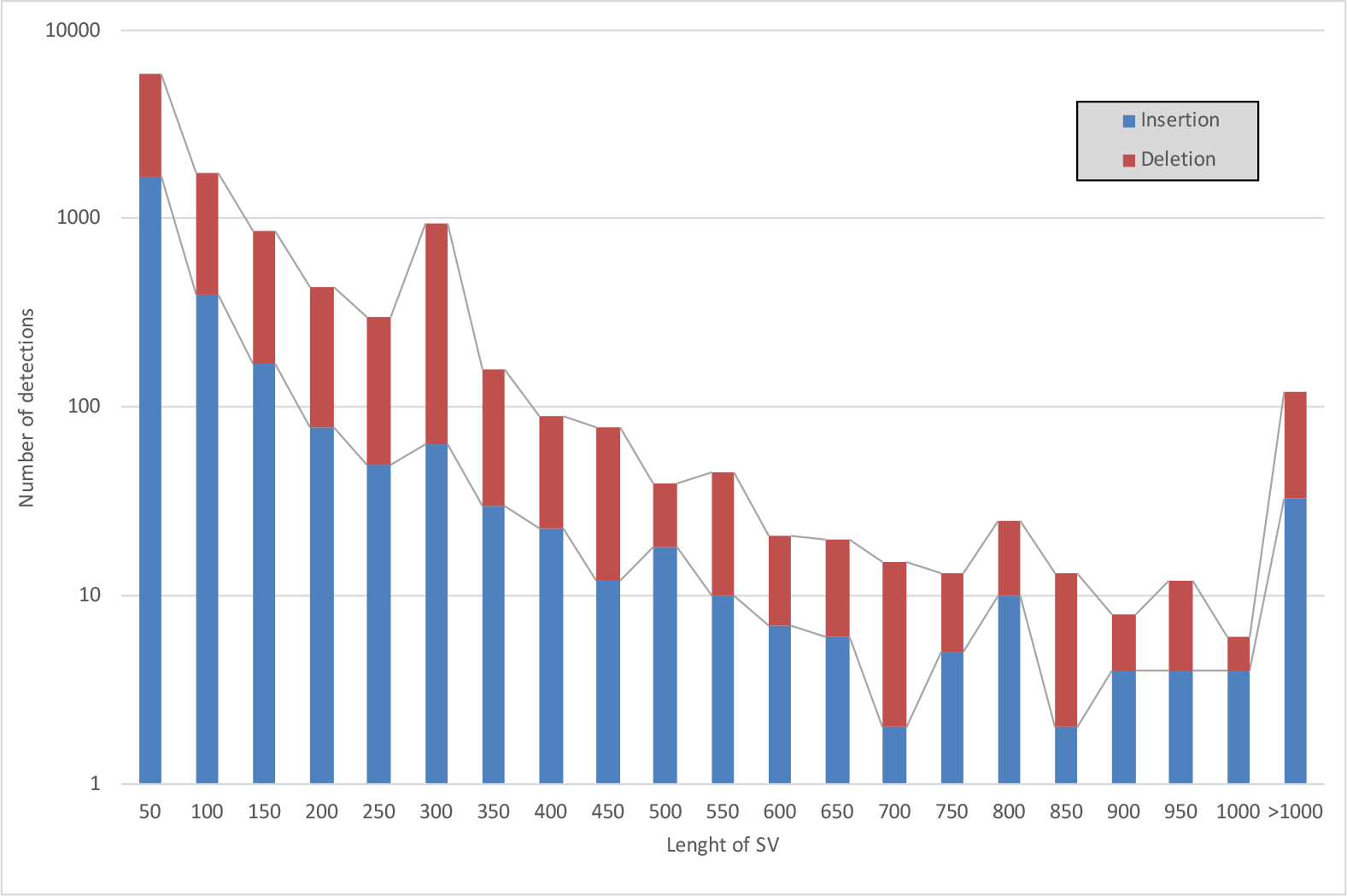
SV length distribution of NA12878. The peak at a length of 251-300 is attributed primarily from Alu elements.

### Comparison against a high-confidence truth set

If ConnectedReads generates a mis-assembled contig, then it should be observed via the CIGAR string after applying any read mapping tools. For example, two unrelated unitigs are assembled together, the contig usually is marked as large soft- or hard-clipping in its CIGAR string. In addition, insertions and deletions in the CIGAR string are the candidates should be investigated in the evaluation process. To evaluate the data correctness of ConnectedReads, the SV dataset collected by svclassify (*7*) is used.

Since insertions are more difficult to be detected by most of existing SV callers than deletions, the 68 high-confidence insertions from svclassify are chosen as the insertion dataset in this paper. In this paper, we develop a naïve Insertion Caller, named CIGAR-based Insertion Caller for ConnectedReads (CCC-INS), which is based on CIGAR string generated by minimap2 with default settings. Additionally, two well-known variant callers are selected for performance evaluation: pbsv for PacBio LRS and FermiKit (*18*) for Illumina SRS. Both FermiKit and ConnectedReads adopt assembly-based approaches to prevent reference bias. The results for the insertion benchmark data are listed in Table 3. PacBio LRS data are usually able to cover the whole region of most SVs, so pbsv achieves a 91.2% detection rate. However, FermiKit detects only 28 insertions since it aims to construct the complete sequence for each insertion. Since most WGS samples are sequenced by SRS and have a coverage of approximately 30X, it is quite difficult to reconstruct the complete sequence (including novel insertions) through any computational approaches. Therefore, we adjusts the naïve insertion caller CCC-INS to relax the criterion of detection in three levels, including completely constructed (CC), partially constructed (PC), and potentially detectable (PD). The CCC-INS^CC+PC^ caller has a strict criterion because it is based on both completely constructed and partially constructed insertions. The CCC-INS^CC+PC+PD^ caller has a lenient criterion because it accommodates all three levels of detection. As shown in Table 3, CCC-INS^CC+PC^ and CCC-INS^CC+PC+PD^ achieve 86.8% and 95.6% detection rates, respectively, indicating that using ConnectedReads is able to achieve the same level of performance as PacBio LRS. Therefore, the above experimental results give us confidence to investigate SVs in population-scale data.

**Table 3.** Long insertion dataset in NA12878 from svclassify

| Method | Detection rate (%) | Number of insertions<br>with complete sequences |
| --- | --- | --- |
| pbsv (PacBio)* | 91.2 | 62 |
| FermiKit | 41.2 | 28 |
| CCC-INS<sup>CC+PC</sup> | 86.8 | 32 |
| CCC-INS<sup>CC+PC+PD</sup> | 95.6 | 32 |

\*The result of pbsv from NIST's Genome in a Bottle (GIAB) project is available at ftp://ftp-trace.ncbi.nlm.nih.gov/giab/ftp/data/NA12878/analysis/PacBio_pbsv_05212019/HG001_GRCh38.pbsv.vcf.gz

Therefore, leveraging ConnectedReads is competent at SV detection, especially for insertions. Then, the next question that we are interested in is how many unique SVs exist in each population. According to Kehr’s findings (in Table S4 of (*9*)), there are 372 SVs in all Icelanders. After removing redundant SVs and merging adjacent SVs, 248 distinct SVs are represented as the second set of SV dataset in this paper. We are eager to know whether these 248 SVs are unique to Icelanders or shared by all populations. Therefore, the three samples from different continents shown in Table 1 are processed by ConnectedReads, and then minimap2 is used to generate alignment records for their ConnectedReads contigs. Then, we manually evaluate whether the 248 SVs exist in the three samples and the results are listed in Table 4. More than 95% of the common SVs in all Icelanders are also found in the three different populations, and only one SV cannot be found in any of the three given samples. It is obvious that most common SVs in Icelanders are not unique to Icelanders. Furthermore, approximately 40 SVs should not be classified as SVs in the given samples because they are composed of several adjacent small variants and incorrectly detected as SVs. More details will be discussed in the next section. It is believed that ConnectedReads provides us with higher resolution to observe SVs than some of SV calling tools.

**Table 4.** The common SV dataset from all Icelanders.

| Sample | Race | Detected |  | Undetected |  |
| --- | --- | --- | --- | --- | --- |
|  |  | SV | Multiple small variants | Not found | Uncovered |
| NA12878 | Caucasian | 204 | 39 | 2 | 3 |
| NA24694 | Asian | 198 | 40 | 2 | 8 |
| NA19240 | African | 205 | 39 | 1 | 3 |

## Discussion

### Accuracy of insertion length

In the above section, ConnectedReads not only was complementary to the resequencing pipeline of SRS in terms of uncovered regions but also was able to benefit SV discovery, especially for long insertions. ConnectedReads and PacBio adopt HG38, but svclassify uses HG19. For comparison, the insertions provided by Spiral Genetics are converted from HG19 to HG38 by using UCSC LiftOver. Then, all of SVs from different source or methods are comparable since they are based on the same genome coordinate. As shown in Table 3, at least 85% of long insertions can be detected by using ConnectedReads followed by minimap2 and CCC-INS, and 32 insertions are completely constructed. As shown in Fig. 4, most of the insertions identified by ConnectedReads are almost the same as those identified by Spiral Genetics and PacBio. 24 of 32 insertions identified from ConnectedReads have identical lengths with PacBio. In addition, the remaining insertions have only slight differences. Therefore, ConnectedReads with SRS can construct complete and accurate insertion sequences as well as PacBio LRS can.

**Fig. 4.**
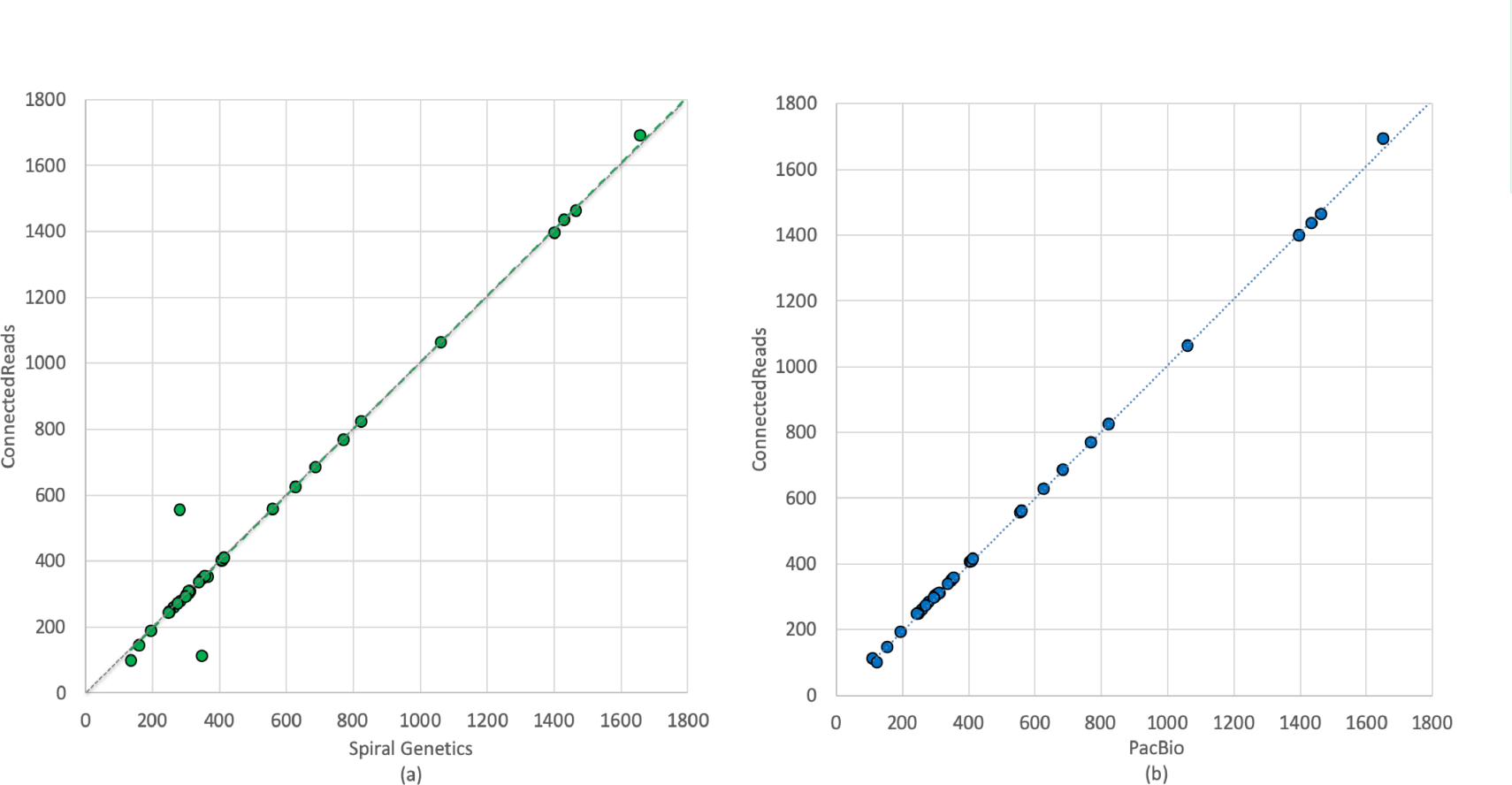
Comparison of insertion lengths among three approaches (genome coordinates are based on hg38). (a) Spiral Genetics vs. ConnectedReads, (b) PacBio vs. ConnectedReads. The gray dashed line is the 1:1 line. The green and blue dashed lines in (a) and (b), respectively, represent the moving average of the comparison.

### Granularity of SV detection

Another interesting phenomenon shown in Table 4 caught our attention. Although common SVs in Icelanders exist in different populations, approximately 40 of the SVs should not be classified as SVs (≥ 50 bps) because they are composed of multiple adjacent small variants, as illustrated in Fig. 5. Fig. 5(a) and Fig. 5(b) contain only 13 SNPs and 13 deletions spanning 60 bps and 80 bps, respectively. When several adjacent variants occur in any individual, most mapping tools have limited information with which to correctly arrange reads with dense adjacent variants and then straightforwardly choose to either employ soft-/hard-clipping or categorize them as unmapped. This limitation will somehow guide most variant callers to identify these regions as SVs. ConnectedReads can prevent such false mapping and help variant callers detect the adjacent variants correctly and precisely.

**Fig. 5.**
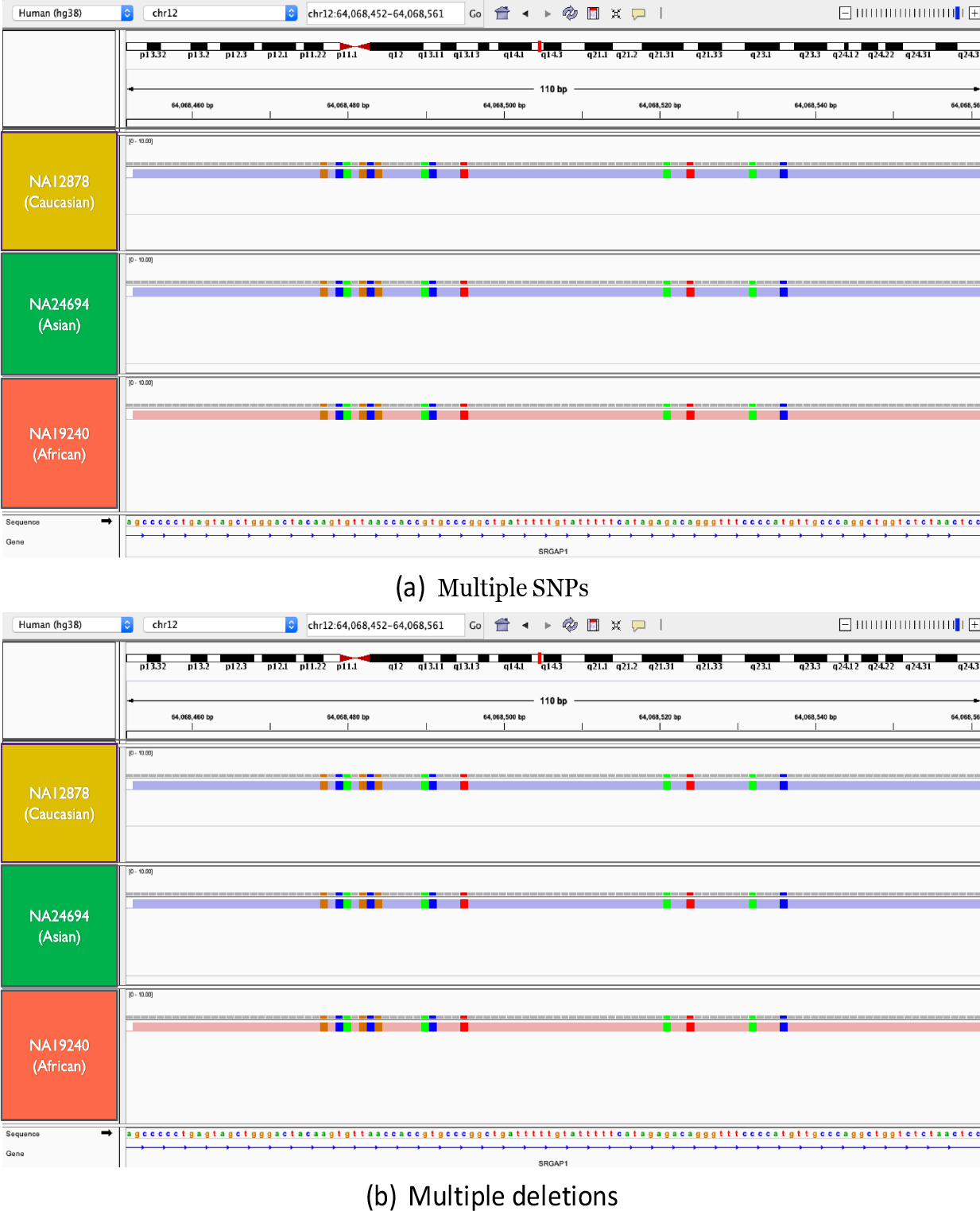
Examples of NRNR sequences (multiple small variants). (a) Thirteen adjacent SNPs in the intron of SRGAP1. (b) Thirteen adjacent deletions in the intron of BICC1.

Furthermore, ConnectedReads is able to mitigate the impact on downstream analysis due to the incomplete human reference genome. The most recent human genome still has many ambiguous areas (N-gaps), and they are mainly located in centromeres and telomeres. Fig. 6 illustrates that two ambiguous gaps can be assembled by using ConnectedReads and that the sequences are totally identical among the three individuals from different populations. This finding gives us strong confidence that most humans might have the same sequence in the two N-gap regions. By randomly selecting two Chinese adults, the occurrence of identical sequences in the N-gap regions is confirmed by Sanger sequencing, as performed by a Clinical Laboratory Improvement Amendments (CLIA)-certificated laboratory. The length of the ambiguous region in Fig. 6(a) should be corrected from 382 to 223 bps. In addition, the length of ambiguous regions in Fig. 6(b) should be shortened from 45 to 31. Based on these cases, leveraging ConnectedReads on cohort-scale WGS dataset is able to provide a cost-effective approach with which to complete the human reference genome.

**Fig. 6.**
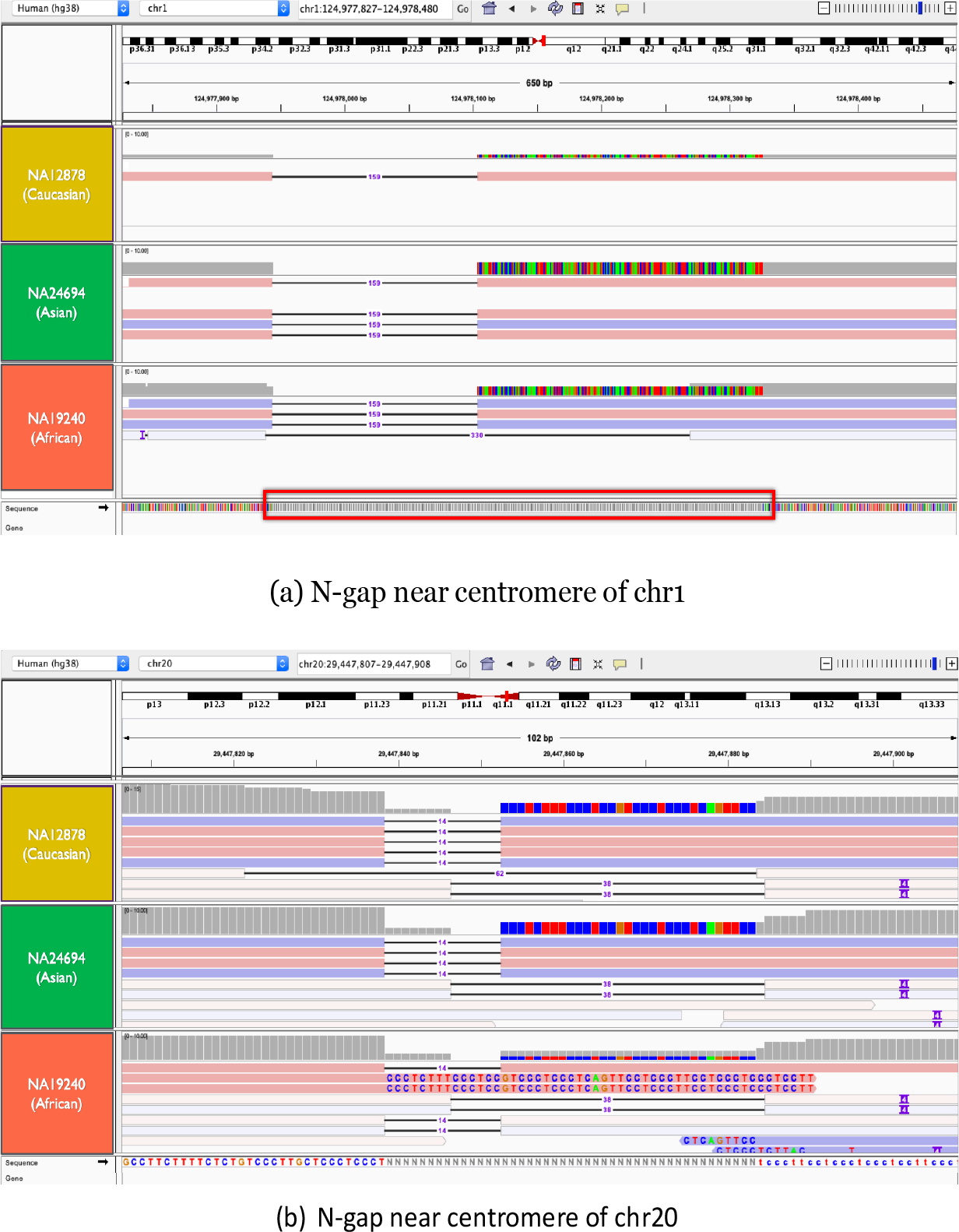
The contigs from three individuals cross the entire N-gap regions near the centromere of chromosome 1 and 20. Based on the alignment result, the sequences in the N-gap are identical among the three individuals. (a) The genomic location of IGV is chr1:124,977,874-124,978,413 of HG38. (b) The genomic location of IGV is chr20:29,447,807-29,447,908 of HG38.

### Correctness of reconstructed contigs

The next topic for the evaluation of ConnectedReads is correctness. To comprehensively investigate the data correctness of contigs constructed by ConnectedReads, all of imperfect matched sequences should be considered because any wrong constructed contigs should not be perfectly aligned to reference genome. Therefore, insertion sequences, soft-clipped sequences and unmapped contigs are selected and then identified as being from *Homo sapiens* or just chimeric DNA sequences resulting from false reconstruction. Take NA12878 for example, there are 37 insertions, 384 soft-clipped and 411 unmapped contigs by removing sequences with a length < 1,000 bps. By using BLAST to find any homologous sequences in the National Center for Biotechnology Information (NCBI) non-redundant sequence (nr) database, each sequence can be identified as human or not. As shown in Fig. 7, 35.1%, 46.6% and 37.7% of the insertions, soft-clipped sequences and unmapped contigs are identical to *Homo sapiens* DNA sequences in the nr database. As the threshold of similarity is continuously lowered, more evidence can be found to support the sequence assembly correctness of ConnectedReads. However, there are six sequences without any support when the threshold is set to 20%. Two of the contigs have low-complexity content, and two are matched to several entries but with less than 10% support. The last two unmapped contigs (CONTIG-8337086 and CONTIG-15793805) are more than 10 Kbps in length. CONTIG-15793805 covers all of CONTIG-8337086 in reverse-complement mode, so CONTIG-15793805 is represented for evaluation. In Fig. 8, CONTIG-15793805 is almost fully covered by several long *Homo sapiens* sequences, and some of these sequences are highly overlapped (at the position of 3000-3500 and 7000-7700). Therefore, the results give us more confident that these non-reference contigs constructed by ConnectedReads should be from *Homo sapiens*. Although more effort on verifying data correctness is emerging and expected, we believe that ConnectedReads achieves high data correctness based on the above findings.

**Fig. 7.**
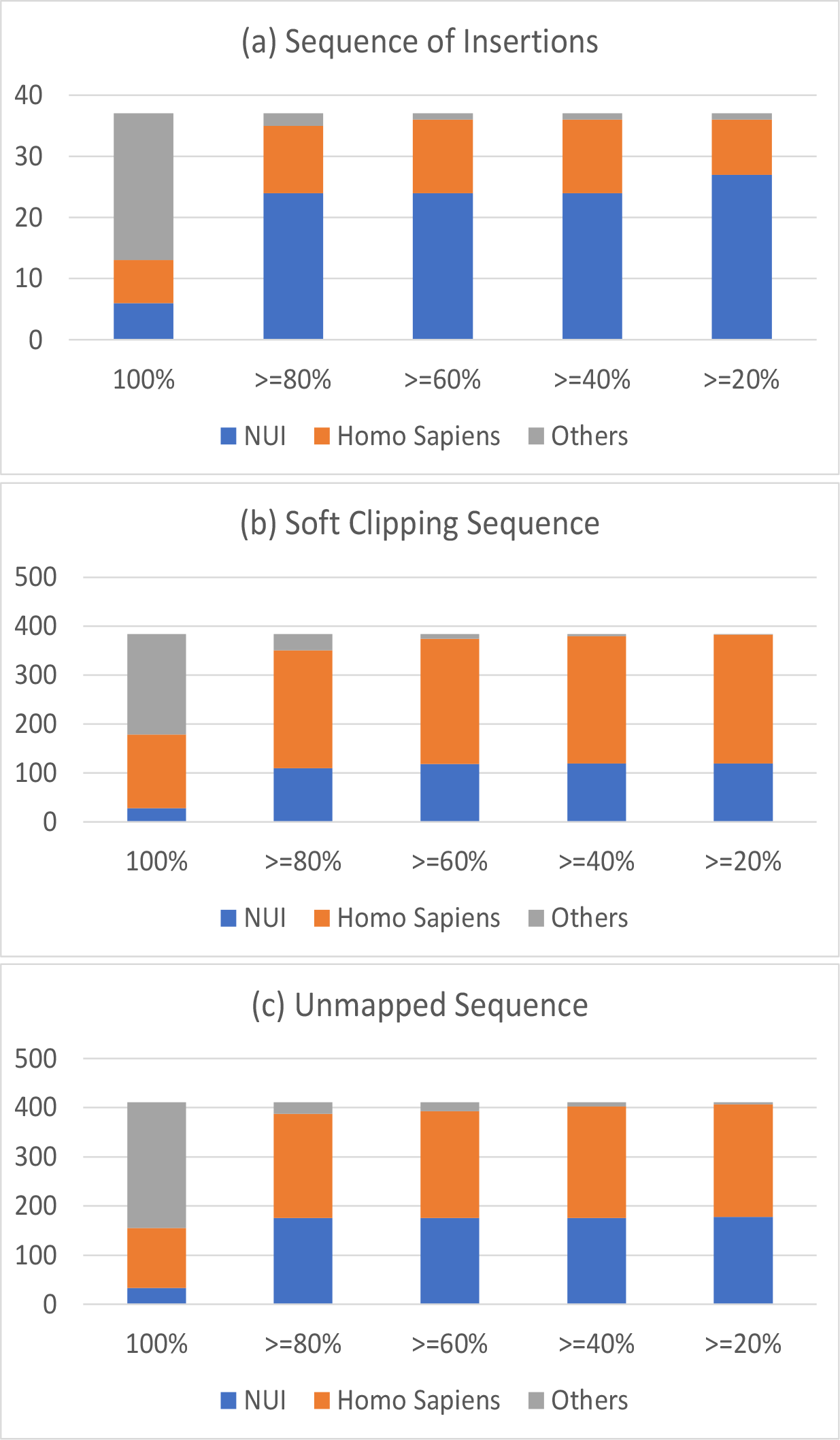
Distribution of BLAST queries for non-reference sequences identified by ConnectedReads. (a) From insertions, (b) from soft-clipped sequences and (c) from unmapped contigs. The x-axis shows the similarity of the query results by BLAST. NUI means that the matched result is annotated as a non-reference unique insertion. *Homo sapiens* means that the matched result is from *Homo sapiens* or Human BAC.

**Fig. 8.**
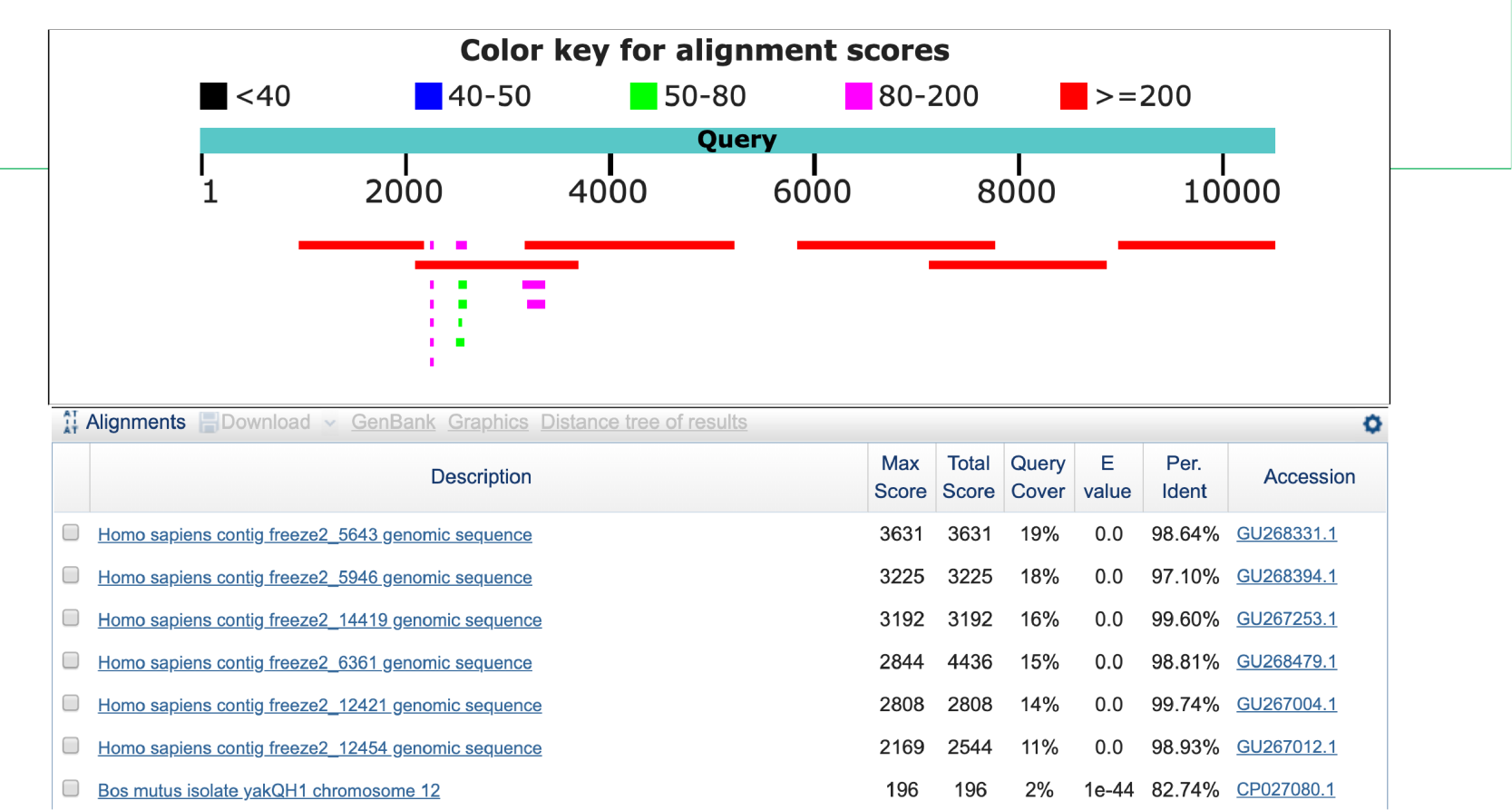
BLAST result for CONTIG-15793805. Six long *Homo sapiens* contigs are identified with low E-values. Since some of them overlap, the alignment result gives us strong confidence in the data correctness of ConnectedReads.

### Translocation-based insertions

Another way to evaluate the data correctness of ConnectedReads is to check whether there is any translocation. If any two unrelated sequences are incorrectly assembled together, it will cause a fake translocation event, in which a contig is mapped to multiple chromosomes. Table 5 lists the 13,686 contigs with multiple alignment records on two chromosomes in sample NA12878.

**Table 5.**
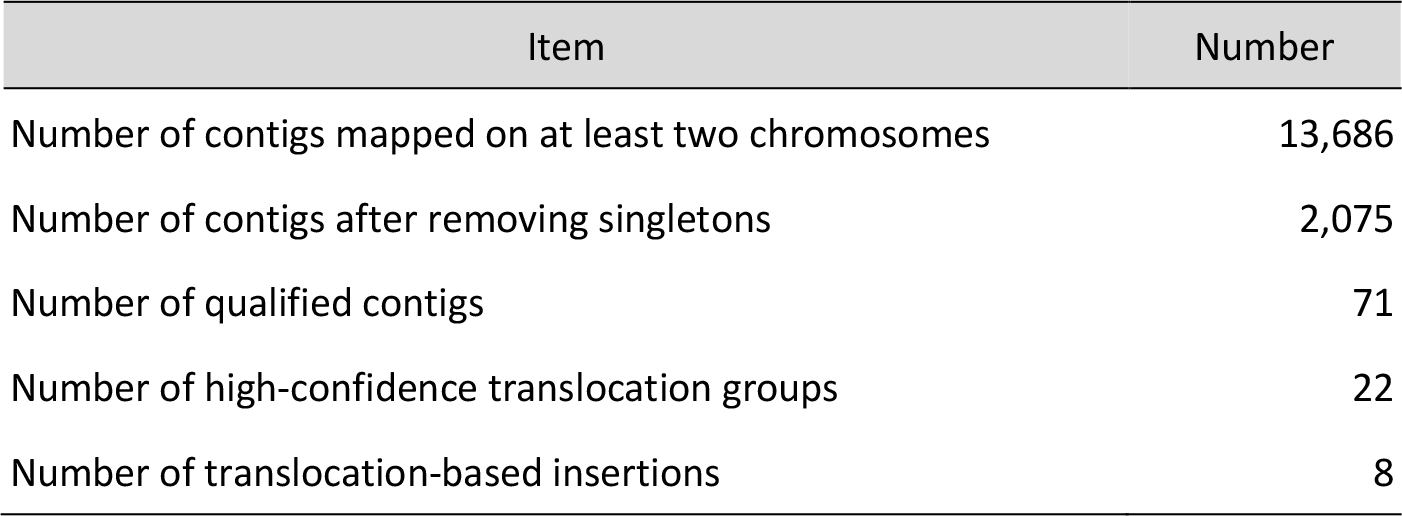
Translocation-based insertions in NA12878.

Generally speaking, a translocation-based insertion should have at least three breakpoints (two for translocation sequences and one for insertion) on genome reference, so it’s difficult to find a single contig with three breakpoints. If a breakpoint for insertion is mapped in only one contig (named as singleton), it won’t be a candidate for translocation-based insertions. After removing singletons and filtering out the contigs with low mapping quality (MAPQ < 60), 71 qualitied contigs are represented by 22 translocation groups.

Interestingly, eight translocation groups have one clear breakpoint on one chromosome but two breakpoints on another chromosome. Those translocation-based insertions can be observed from the VCF of NA12878 generated by pbsv. As shown in Fig. 9(a), for example, CONTIG-6374680 and CONTIG-1880453 have identical breakpoints at chr3:110694547. However, CONTIG-6374680 and CONTIG-1880453 have their own breakpoints at chr1:108952231 and chr1:108952660, respectively. Thus, in NA12878, the 430-bp sequence in intron 1 of CLCC1 is inserted at chr3:110694547. Fig. 9(b) also shows that the 697-bp sequence in exon 10 of BTBD7 is inserted into intron 1 of SLC2A5. These translocation groups are called translocation-based insertions. ConnectedReads proposes a naïve way to investigate these translocation-based insertions. In summary, ConnectedReads not only constructs genome sequences precisely but also facilitates SV detection by mitigating reference mapping bias.

**Fig. 9.**
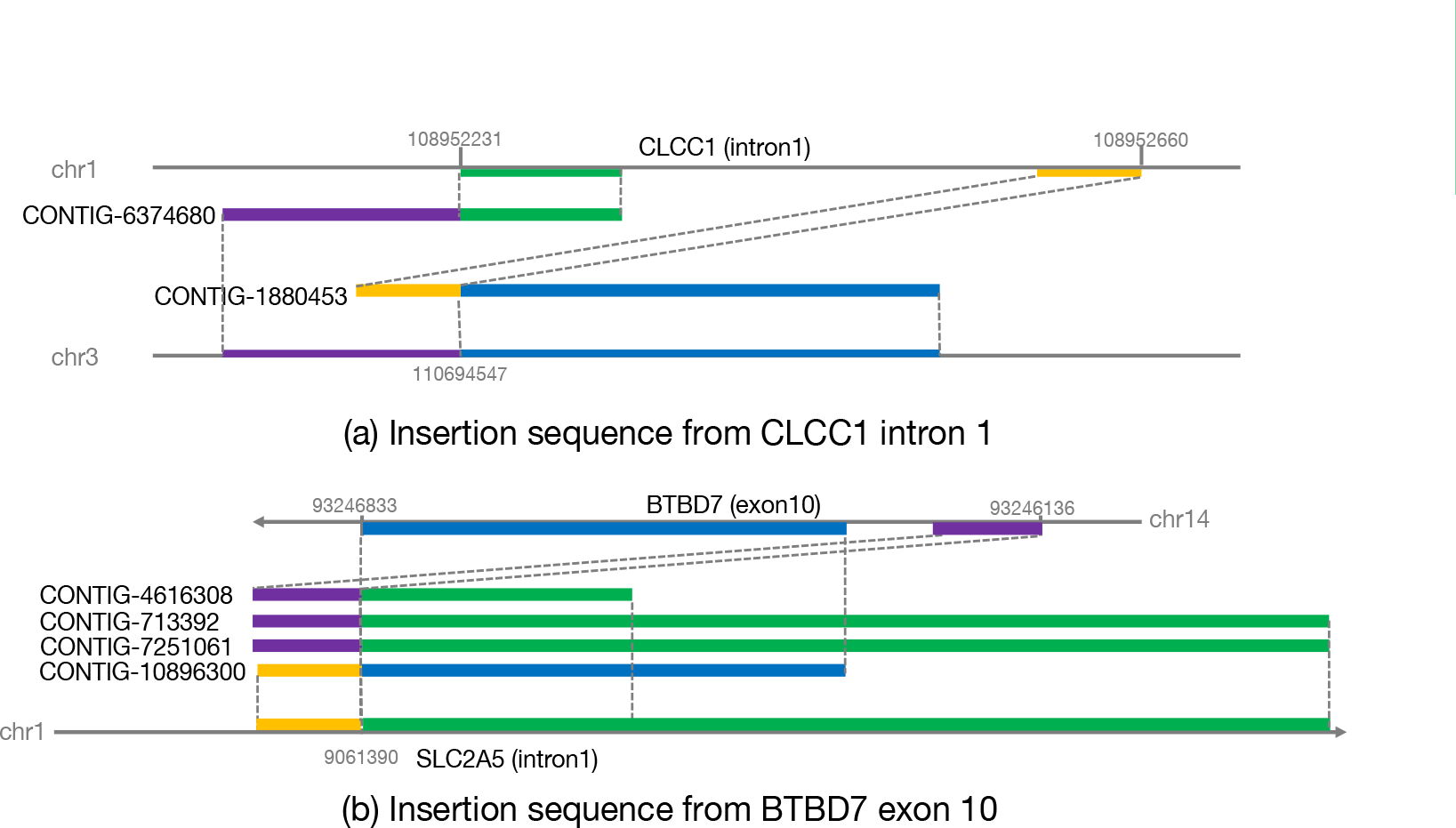
Examples of translocation-based insertions. (a) The 429-bp sequence from intron 1 of CLCC1 is inserted at chr3:110,694,547. (b) The 697-bp sequence from exon 10 of BTBD7 is inserted into intron 1 of SLC2A5.

## Conclusions

NGS data analysis remain a considerable challenge due to an incomplete reference genome, mapping errors and high sequence divergence. By leveraging ConnectedReads, SRS is able to be constructed from short reads to long assembled contigs. Using ConnectedReads not only prevents mapping errors but also facilitates SV discovery. In addition, ConnectedReads are able to mitigate the influence on reference incompleteness. In summary, ConnectedReads can serve as an NGS gateway for facilitating downstream data analysis, such as false positive prevention, SV detection, and haplotype identification.

## Materials and methods

### Workflow

ConnectedReads leverages Apache Spark (*6*), a distributed in-memory computing framework, to accelerate its whole workflow, as illustrated in Fig. 10. To fully utilize the power of the distributed framework, the preprocessing step involves splitting a large compressed file into several small files and then uploading those files into the Hadoop distributed file system (HDFS) since most of the WGS samples exist in two separate FASTQ files in GZIP format. First, we adopt Apache Adam (*19*), as shown in Fig. 10(a), to transform data from FASTQ format to column-based Parquet format for data parallel access. To facilitate data processing in the following steps, we extend Adam to not only encode paired-end information and barcodes into the read name column but also place all reads in different subfolders based on their sequencing quality and sequence complexity. Then, we propose a distributed suffix tree algorithm with supervised graph mining on Spark to mitigate the influence of improper string graph construction due to sequencing errors. Using an outlier detection algorithm on a suffix tree, the process in Fig. 10(b) can be configured as a highly sensitive error detector for low-coverage regions. After that, we are able to adopt the parallel string graph construction shown in Fig. 10(c) to represent the relation of each qualified read and the read overlaps by suffix-prefix information. More details are available in our previous paper (*20*). Based on the string graph, we propose the parallel haplotype-sensitive assembly (HSA) depicted in Fig. 10(d) to construct assembled contigs; the detailed procedure of this module will be described in the next session. For example, these generated contigs (Fig. 10) include (i) heterozygous SNPs, (ii) heterozygous deletions, (iii) heterozygous insertions and (iv) homozygous SNPs. In summary, ConnectedReads transforms noisy short reads with low quality and sequencing errors to long assembled and qualified contigs.

**Fig. 10.**
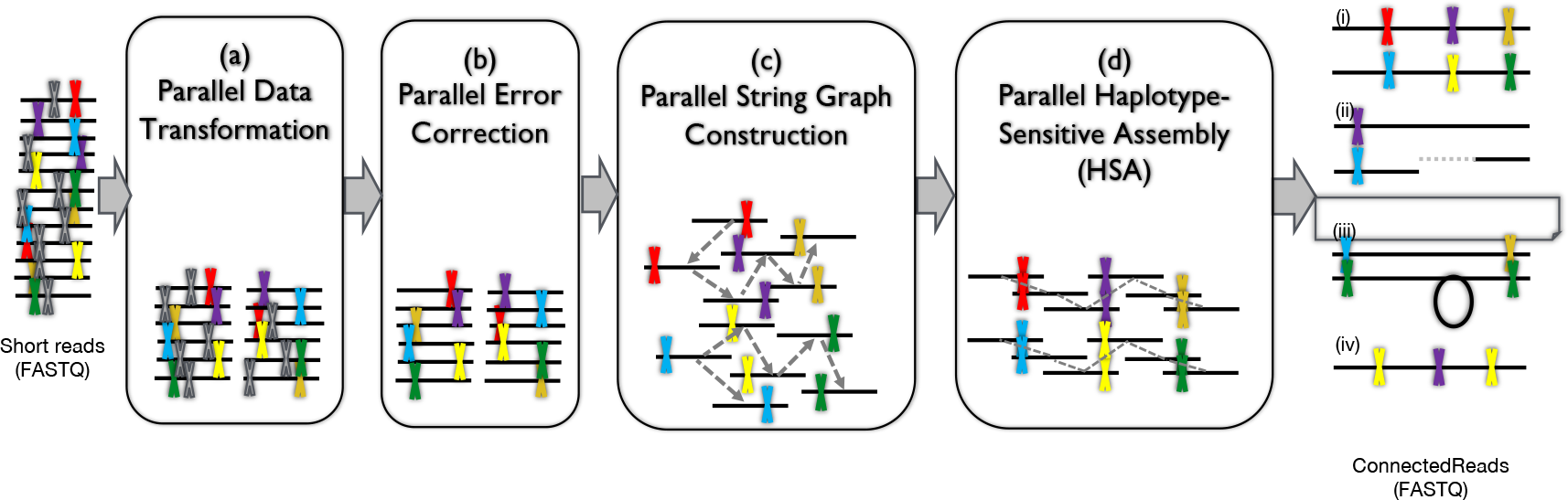
Workflow of ConnectedReads. (a) Parallel data transformation. (b) Parallel error correction. (c) Parallel string graph construction. (d) Parallel haplotype-sensitive assembly (HSA).

### Parallel HSA

ConnectedReads adopts a string graph, a lossless data representation that is fundamental for many de novo assemblers based on the overlap-layout-consensus paradigm (*21, 22*), to represent the overlaps of each read. Here, we propose a Spark-based HSA based on a string graph. By leveraging GraphFrame, HSA is able to perform efficient and scalable graph traversal operations and supervised graph mining algorithms on Apache Spark. Before introducing the detailed algorithms of HSA, we formally define the notations of the sequencing data and string graph that we will use in the following sections.

### Definitions and notation

Let *G(V, E)* be a directed graph and *S*, *T* ⊆ *V*. We define *V* = {*v_1_*,*v_2_*, …,*v_k_*} and *E(S, T)* to be the set of edges going from *S* to *T*, i.e.,

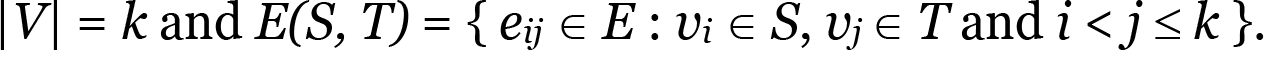

#### Definition 1.

Let *R* be the set of short reads and *R^RC^* be the reverse complement set of *R. G* and *G^RC^* are the string graphs based on *R* and *R^RC^*, respectively. If *e_ij_* exists in *G* and *i, j ϵ R*, then *e_mn_* also exists in *G^RC^* and *m,n ϵ R^RC^* such that *m* and *n* are the reverse complements of *j* and *i*, respectively.

We define *G^EXT^(V, E)* to be the expanded graph of vertices *V ϵ R ∪ R^RC^* and edges *E* to be the set of edges in either *G* or *G^RC^*.

#### Definition 2.

Let *IN*(*v_i_*) and *OUT*(*v_i_*) be the number of in-degree and out-degree edges of *v_i_*, respectively. We define a vertex *v_i_* to be

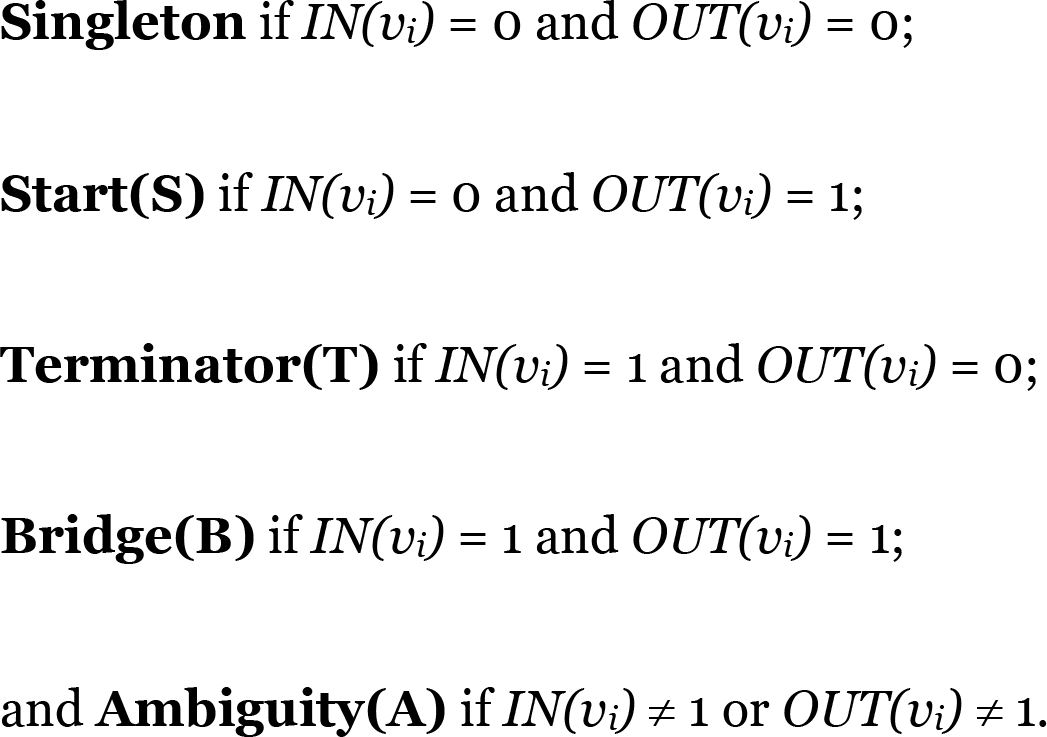

#### Definition 3.

We define the priority order of the vertex labels to be **T** > **S** > **B** and **A**.

This means that **B** could be relabeled as **S** or **T**. Once a vertex becomes **T**, it will always be **T** regardless of what the graph property propagation is.

To keep the depth information in FASTQ format, a new quality encoding function is proposed.

#### Definition 4.

Let *L* and *N* be the numbers of layers for quality and depth, respectively. *L*N* should be 42 if Phred33 is adopted. Let *Q[i]* and *D[i]* be the quality and depth of the *i*-th base of the given contig, respectively. We define the quality encoding function *EncoderQ[i]* and the depth encoding function *Encoder_D_[i]* to be

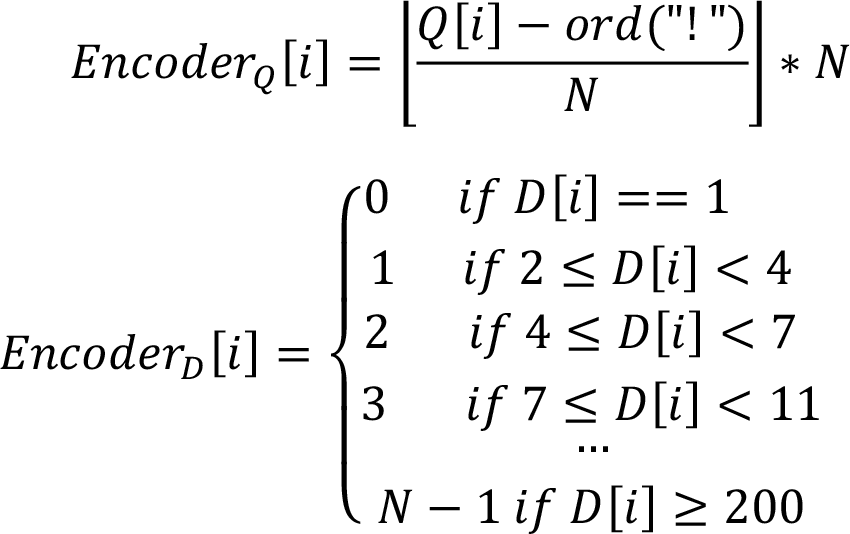

In addition, we define the quality-depth (QD)-encoding function *Encoder_QD_[i]* to be

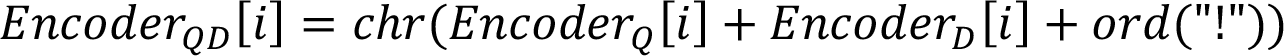

### System flow

In Fig. 11, HSA makes use of five modules, namely, (a) string graph expansion, vertex connectivity classification, (c) the Start-Bridge-Terminator (S-B-T) traversal algorithm, (d) the ploidy-based routing algorithm and (e) the triplet traversal algorithm. To simplify the following graph traversal algorithms, each vertex and its reverse complement should be separated in a string graph. This means that the string graph *G* generated by Fig. 10(c) should first be expanded by Definition 1 and named *G^EXT^*, which contains *G* and *G^RC^*. For that reason, Fig. 11(a) generates the expanded string graph with all reads and their reverse complements. After removing all singletons in *G^EXT^*, the remaining vertices will be classified by the vertex connectivity classification module shown in Fig. 11(b) into four classes, which are defined in Definition 2: start (**S**), bridge (**B**), terminator (**T**) and ambiguity (**A**). To generate assembled contigs, the graph traversal/pairing algorithms shown in Fig. 11(c-e) are proposed based on the above properties of vertices and described comprehensively in the next section.

**Fig. 11.**
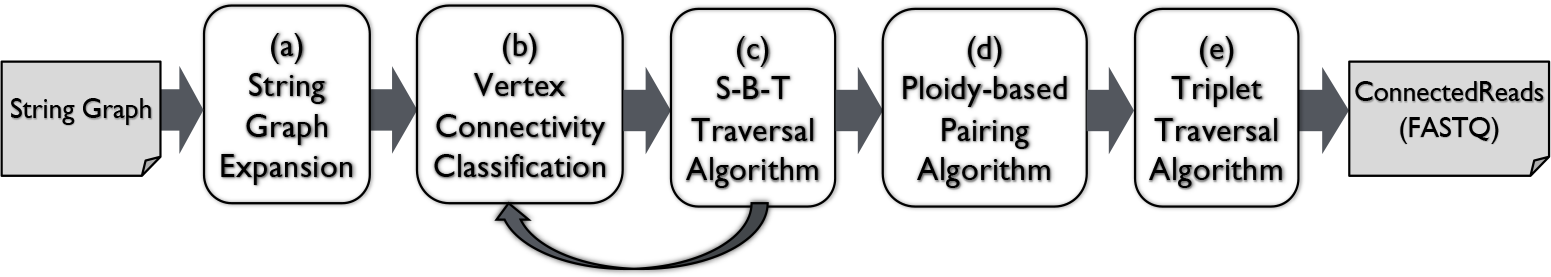
Parallel haplotype-sensitive assembly (HSA). (a) String graph expansion. (b) Vertex connectivity classification. (c) S-B-T traversal algorithm. (d) Ploidy-based paired algorithm. (e) Triplet traversal algorithm.

### Graph mining algorithms

ConnectedReads leverages Apache Spark and its derived packages to propose three efficient and scalable graph algorithms for large graphical datasets, i.e., string graphs. Spark GraphFrame is a powerful tool for performing distributed computations with large graphical datasets. In addition, Spark Dataset is a type-safe interface that provides the benefits of resilient distributed datasets (RDDs) and Spark SQL optimization. By leveraging GraphFrame and Dataset, we propose several Spark-based graph traversal/routing algorithms for HSA with high performance and scalability.

From our observations of NGS sequencing data, two challenges must be overcome if we want to enhance the performance of graph operation for HSA. The first challenge in the string graph involves long diameters, and the second challenge is how to properly connect the vertices with multiple in-/out-degrees. Taking NA12878 as an example, more than 90% of the vertices are **B,** and the longest diameter from **S** to **T** is more than 500. This means that the traversal algorithm must perform the propagation operation in at least 500 iterations from one vertex (**S**) to another vertex (**T**). For most graph frameworks, the performance of a graph algorithm is strongly related to the number of iterations for its traversal operation. Here, we propose the S-B-T traversal algorithm, shown in Fig. 11(c), to connect all vertices from **S** to **T** via all adjacent **B**s. By following Definition 3, the data preprocessing flow for the expanded string graph is applied, as illustrated in Fig. 12. First, all vertices before and after any vertex **A** should be relabeled as **T** and **S**, respectively. To overcome the long-diameter problem, the mechanism used to relabel **B** to **T** by customized random selection is applied to shorten the diameter of the given graph, and then, the terminator-intensive propagated string graph is acquired. Based on the terminator-intensive graph, the S-B-T traversal algorithm, which integrates a belief propagation algorithm with iterative graph merging, is able to theoretically reduce the time complexity from O(*N*) to 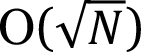 (*N*: the number of iterations for graph propagation).

**Fig. 12.**
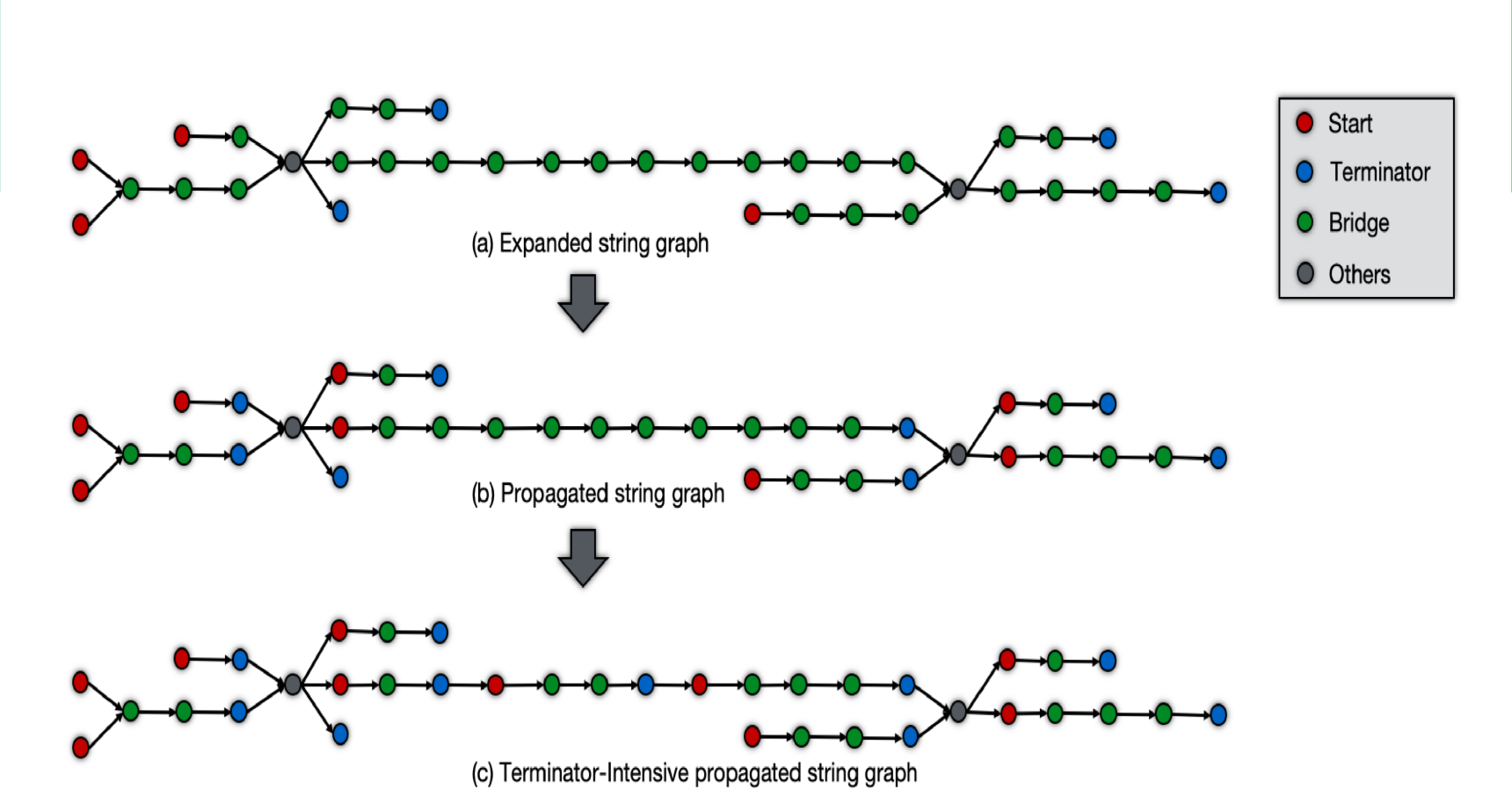
An example to demonstrate the graph preprocessing workflow for the S-B-T traversal algorithm. (a) Expanded string graph. (b) Propagated string graph. (c) Terminator-intensive propagated string graph.

When each simple routing path (i.e., **S** to **T** via **B**s) is completely traversed and merged into a single vertex in the new graph, the majority of the vertices have multiple in-/out-degrees. Therefore, we have to solve the second challenge - how to select a proper routing path in the multiple-in-/out-degree graph. Many useful indicators enable us to perform correct routing, such as read pairs, barcodes and read overlaps. The pseudo code of the ploidy-based pairing algorithm shown in Fig. 11(d) is as follows:

~~~
def **ploidy_based_pairing**(G: GraphFrame for string graph, N: int for ploidy) {
   Candidates = ∅
   V_a_ = all of **A**s in G
   for each v ∈ V_a_ {
       I_v_ = the set of vertices point to v
       O_v_ = the set of vertices pointed by v
       D_v_ = **de_noise**(I_v_, v)
       T = ∅
       for each u ∈ D_v_ {
         (t, s) = **find_best_matching**(u, v, O_v_)    #t: triple; s: score
if s ≥ MIN_THRESHOLD then
   add (t, s) into T
}
sort T by score
add the top N triples from T into Candidates
}
return Candidates
}
~~~

To mitigate false assembly due to sequencing errors, the function **de_noise()** is used to remove the noisy vertices in *Iv* by using a naïve clustering algorithm based on the read-pair information in this paper. In addition, the function **find_best_matching()** adopts a tripartite clustering algorithm based primarily on the number of supports from read-pair information in (*u*, *v*), (*u*, *O_v_*) and (*v*, *O_v_*) to find the best combination for sequence assembly. Using the ploidy-based pairing algorithm, we obtain sufficient information to overcome the second challenge and then apply the triplet traversal algorithm shown in Fig. 11(e) to construct the assembled contigs by traversing the aggregated graph from each **S** to **T** via the triple set generated by the ploidy-based pairing algorithm. The triplet traversal algorithm involves almost the same procedures as the S-B-T traversal algorithm, except for the linkage of vertexes. In summary, HSA leverages Apache Spark to efficiently generate assembled contigs from a large string graph.

### Performance

Here, sample NA12878 is processed by ConnectedReads on our Spark cluster. It takes approximately 18 hours for the 30X WGS sample; the detailed performance of ConnectedReads is described in Table 6.

**Table 6.** Execution time for NA12878

| Module | Time (hours) | # Executors | # Cores per executor |
| --- | --- | --- | --- |
| (a) Parallel data transformation | 1.9 | 9 | 17 |
| (b) Parallel error correction | 8.1 | 15 | 8 |
| (c) Parallel string graph construction | 4.3 | 9 | 17 |
| (d) Parallel haplotype-sensitive assembly | 4.0 | 9 | 17 |
| Data export from HDFS to local disk | 0.05 | 9 | 4 |
| Total | 18.3 |  |  |

\*There are 9 computing nodes (each node has two E502650 v4 with 512 GB of memory)

### QD encoder/decoder

Since ConnectedReads not only connects reads with their overlaps but also aggregates identical reads, the size of ConnectedReads contigs will be reduced by at least 90% in comparison with that of contigs from short-read data. However, the downstream analysis tools might not work well due to the loss of depth information. We have to take the depth information into consideration and keep the output compatible with FASTQ format. Therefore, the QD encoder and decoder are proposed based on Definition 4 to mitigate the impact on information loss. The mechanism of the QD-encoding function is quite flexible, allowing it to fit most use cases. For example, if the depth information is more critical than quality, (L, N) = (3, 14) is the best choice. Fig. 13 provides an example to demonstrate how the QD encoder transforms short reads into ConnectedReads contigs. The data reduction rate is approximately 77% (from 144 characters to 33 characters in both sequence and quality). In addition, Fig. 14 shows the result of applying the QD-decoding function. The recovery rate in terms of sequences and quality is 95.8% (138/144) and 47.3% (71/144), respectively. Therefore, ConnectedReads provides an efficient QD-decoding tool for some use cases that heavily leverages depth information.

**Fig. 13.**
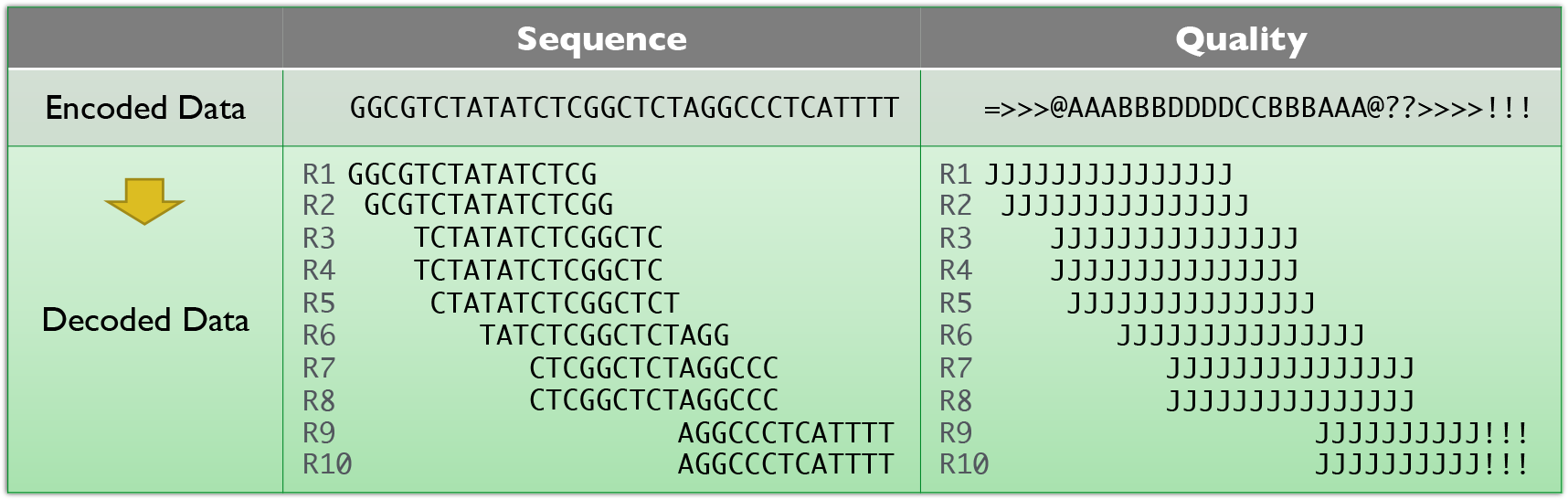
An example application of the QD encoder (taking (L, N) = (3, 14) as an example).

**Fig. 14.**
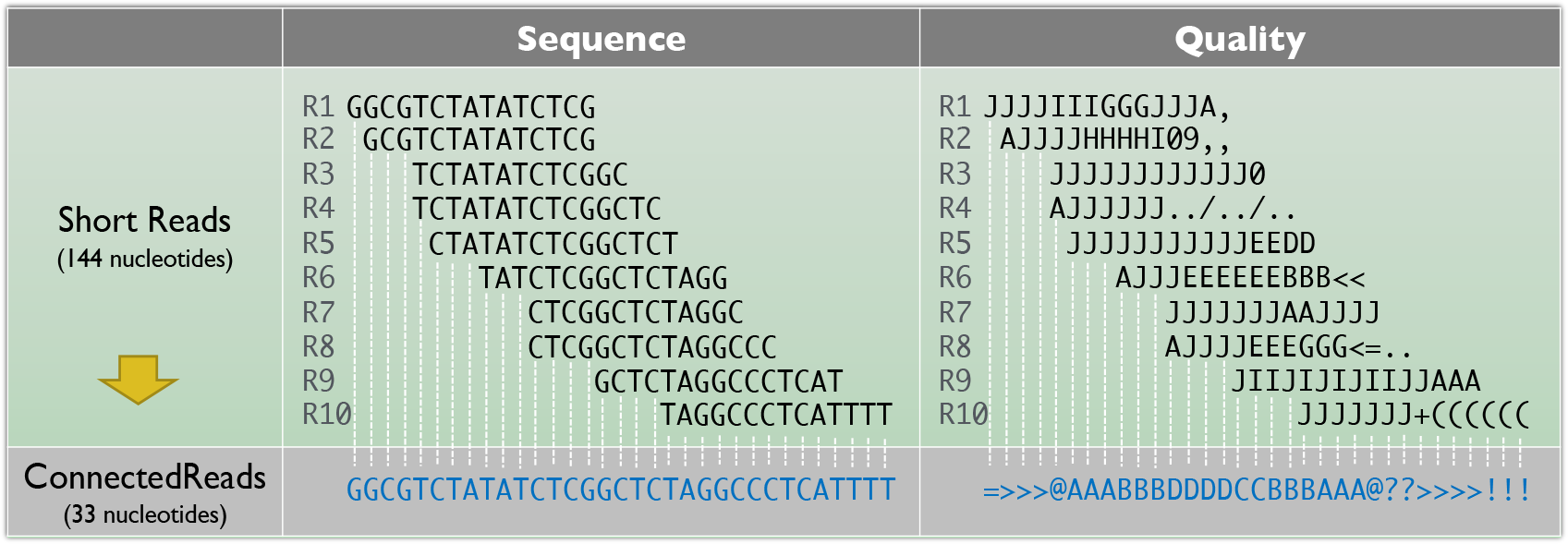
An example application of the QD decoder (taking (L, N) = (3, 14) and read length = 15 as an example).

### Availability of data and materials

Data: The raw sequencing data discussed in this manuscript are deposited on the European Bioinformatics Institute (EBI) and NCBI websites. NA12878 is available from https://www.ebi.ac.uk/ena/data/view/ERR2438055; NA24694 and NA19240 are available from https://www.ncbi.nlm.nih.gov/sra/SRX1388455 and https://www.ncbi.nlm.nih.gov/sra/SRX4637790, respectively.

Codes: The source code and scripts for the ConnectedReads workflow and the related experiments discussed in this paper are available at https://github.com/atgenomix/connectedreads.

## Supporting information

N-Gap validation

insertion sequence

soft-clipping sequence

unmapped sequence

## Acknowledgements

We are grateful to Prof. Chen Chia-Hsiang and his team for supporting the Sanger sequencing validation. In addition, Cheng Yun-Chian and Prof. Chen Chien-Yu assisted in algorithm design in the early stage. The results presented in the paper are based on the data generated by the Atgenomix Seqslab Cloud platform.

## Contributions

CS, SW, YL and MC developed the algorithms and implemented the tools for data transformation and error correction. CS developed the algorithms and implemented the tools for string graph construction. CS and SW developed the algorithms and implemented the tools for haplotype-sensitive assembly. CS collected the sequencing data and carried out the benchmarks based on the sequencing dataset. CS carried out the experiments, developed the structural variant caller and carried out the related experiments. CS prepared the manuscript with input from all other authors. All authors read and approved the final manuscript.

## Ethics declarations

Ethics approval and consent to participate

This article is a preprint version and can be accessed on https://www.biorxiv.org/content/10.1101/776807v1. This article is not published nor is under publication elsewhere.

## Consent for publication

Not applicable.

