## Supplementary material for "ConnectedReads: machine-learning optimized long-range genome analysis workflow for next-generation sequencing": N-Gap validation

### N-Gap case 1

##### Position (Reference: HG38)

chr1:124,977,271-124,978,929

##### Sequence (N-Gap is marked in red)

ATCGCTCGAAATATCCACTTGCAGATCCTACAACGAGACTGTTTCAAAACAGCTCTATCAACAGGATTGTTCAACTCTGTGAGGTGTATGCAGACATCACAAACAAGTTCCTGAGAATGCTTCTGTCTAGTTTTTCTGTGAGGATATTTCCTTTTCCGACATAGGCTTCAAATCGCTAGAAATATCCAATTGCAGATTCTACAAATAGACTGTTTCAAAACTGCTCTCTCAAAGGAAGGTTCAACTCTGTGTGTACAATGCATACATCATAAAGAAGTTTCTGAGAATGCTTCTGTCTAGATTATATGTGAAGATGTTTCCGTTTCCAACATAGGCCTCCAAGCACTCCAAATGAATACTTGCAGATCCTAGAAAAAGAGTGTTTCAAAGTTGCTCTTTCTAAAGAAGTGTTCAACTCTCTGAGTTGAATTCACACATCACAAAGCAGTTTCTCAGCATGCTTCTGTCTAGTTTTTATTTGAAGTTATCTCGTTTCCAAAGAAATCCTAAAACAGCTCCAAATATACACAAGCAGATTCTACAAAAGTAGTGTTTCAGTACTGCTCTATCAAAAGAAATGTTCAACTCTGTGAGTTGAATGCACACATAATAAAGCAGTTTCTGAGAATGCTTCTGTTAAGCTTTTATGTGAAGATATTTCCTTTTCCACCATATGCCGGAAATTGCTCCAAGCATCCACCTACATATCCTGCAAAGAAACTGTTTCTAAACAGCTCTCTCAAGAGGAAGGTGCAGCTCTGTGAGTTGAATGCACACATCACAAACAAATTCCTGAGAATGCTTCTGTCTAGTTTTTCTGTGAAGATATTTCCTTTTCCAACATAGGCTTCAAATCGCTCCAAATATCCAATTGCAGATTCTGCAAAAAGATTGTTTCAAAACTGCTCCCTCAATAGCAAGGTTCAACTCCGTTGGTTGACTTTATACATCACAAAGAAGTTTCTGAGAATGCTTCTGCTTTGTTTTTATGTGAAGATGTTTACTTTTCCACCATAGGCCTCAAAGAACTCCAAACGAACACTTGCAGATCCAAGAAAAAGAGTATTTCAAAACTGCTCTTTCTAAAGAAGTGTTCAACTCTCTGAGTTGAATTCACACATCACAAAGCAGTTTCTGAGAATGCTTCTGTCTAGTTTTTATTTGAAGATAATTCGTTTCCAGCAAAATACACAAAGAACTCCAAATATCCACAAGCAGATTCCACAAAATTAGTGTTTCAGTACTGCGGTATCAAAAGAAATGTTCAGCTCTGTGAGTTGAATGCTCACATCACAAACAAGTTCCTGATAATGCTTCTGTCTACTTTTACTGTGAAGATATTTCCTTTTCCTACATAGGCTTCAAATCGCTCCAAATATCCACTTGCAGATTCTACAAAAAGACTGTTTCAAAACCGCTCTCTCAAAAGGAGGGATCAACTCTGTGAGTTGAATACGTACATCACAAAGTAGTTTCTGAGAAACCTTCTGCCTAGTTTTT

##### Genome Browser

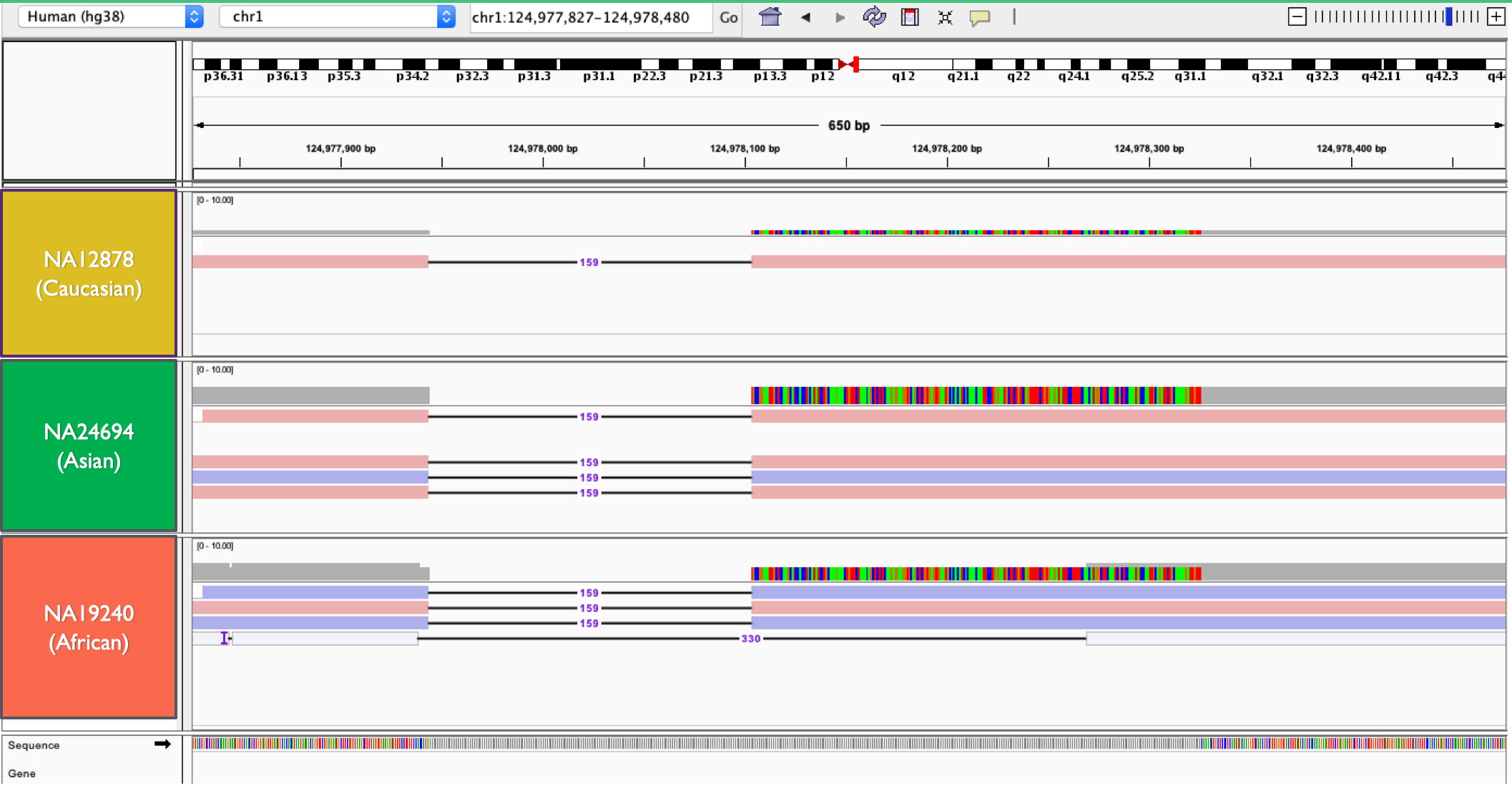

##### PCR Primer

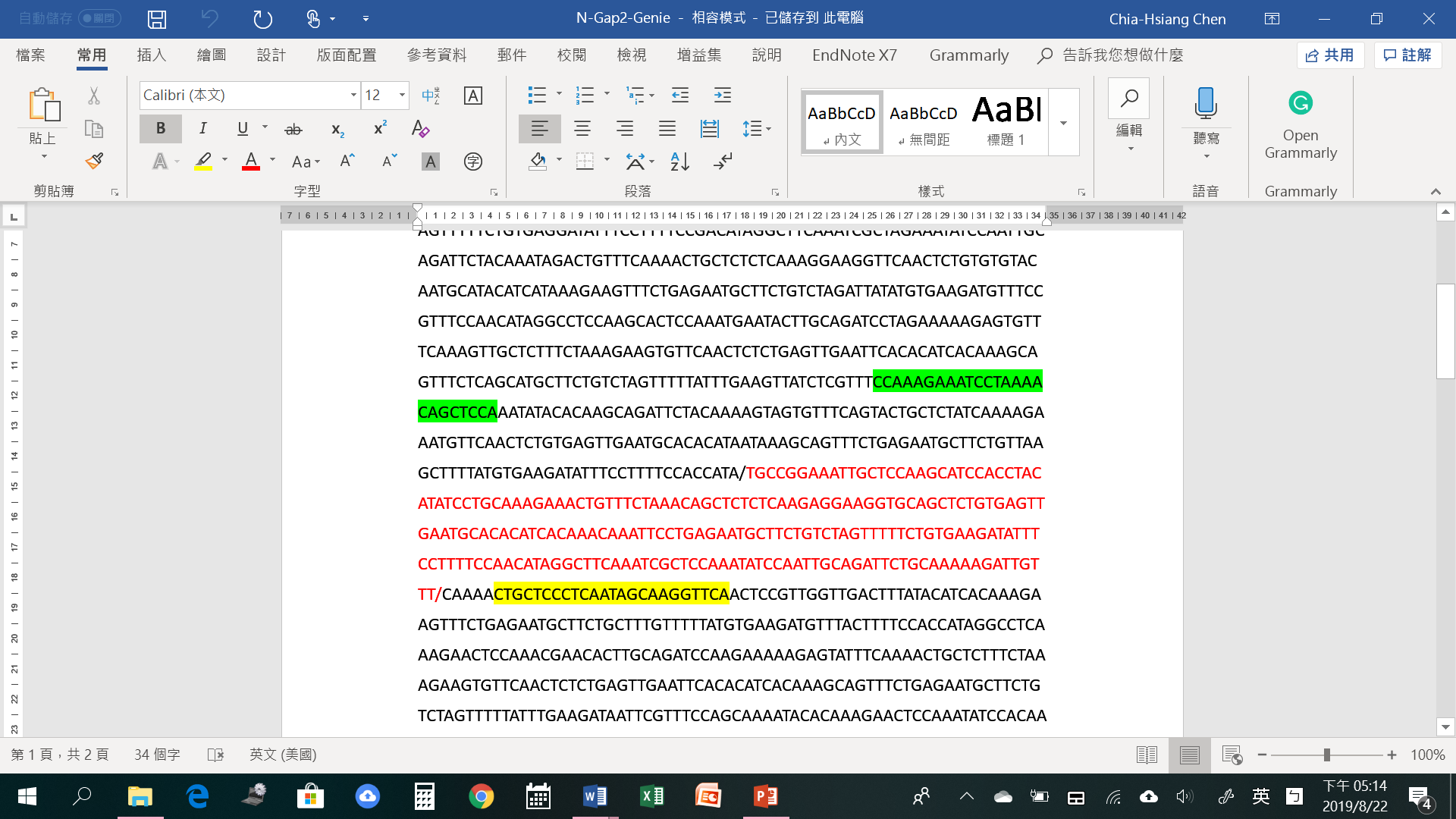

N-Gap2-F: 5’-CCA AAG AAA TCC TAA AAC AGC TCC A-3’ (25 mer)

N-Gap2-R: 5’-TGA ACC TTG CTA TTG AGG GAG CAG-3’(24mer)

408 bp

##### PCR Result

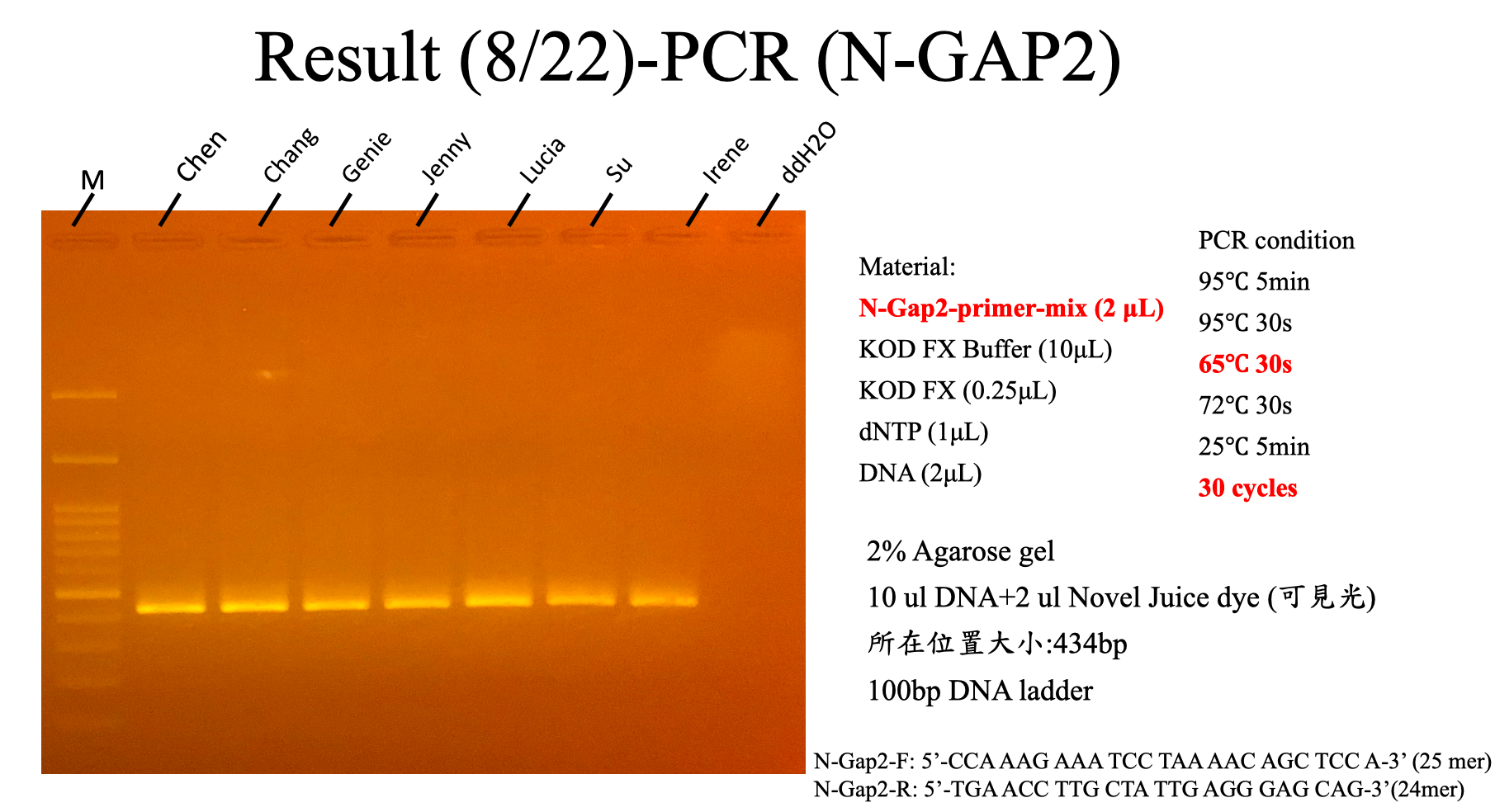

##### Sanger Validation

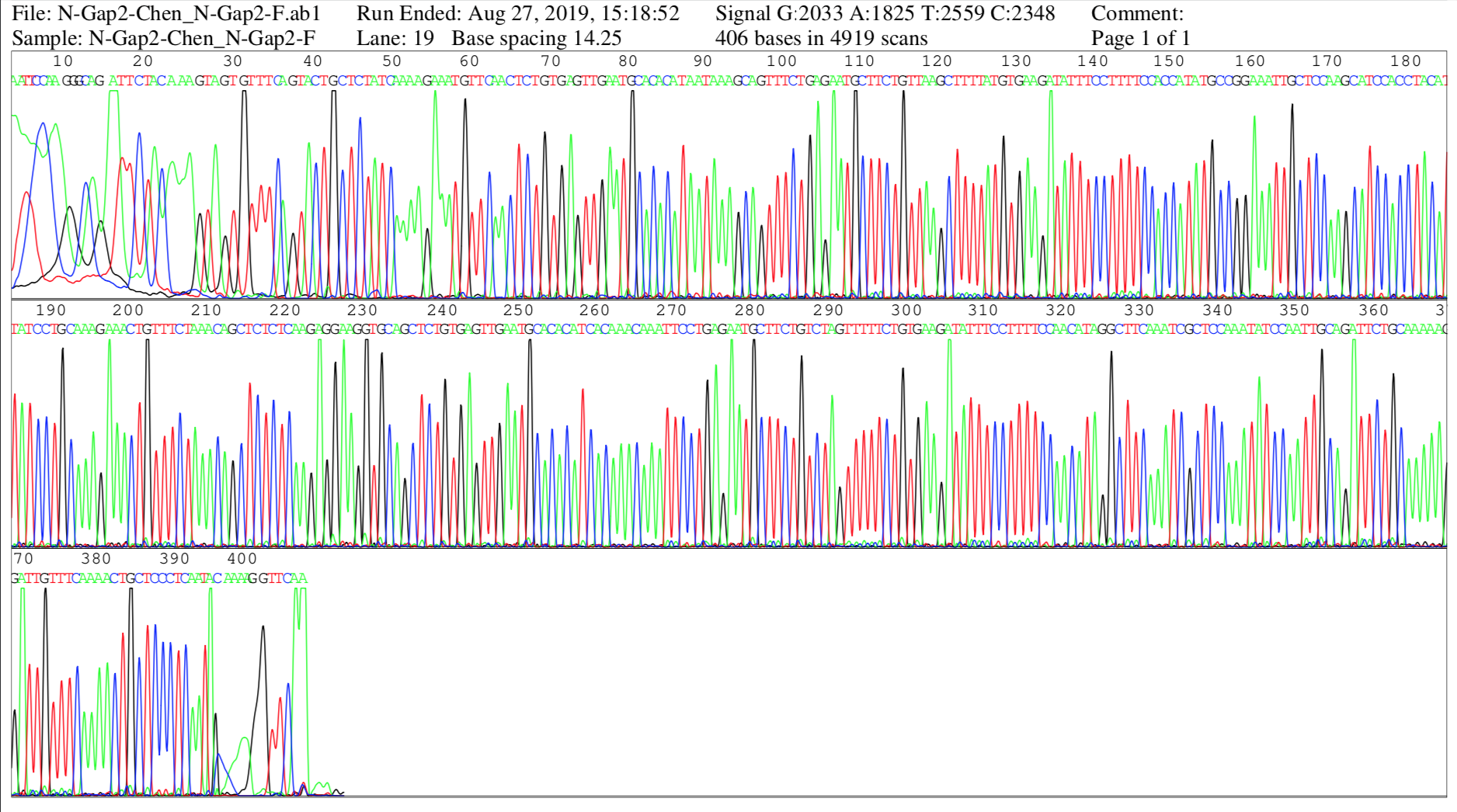

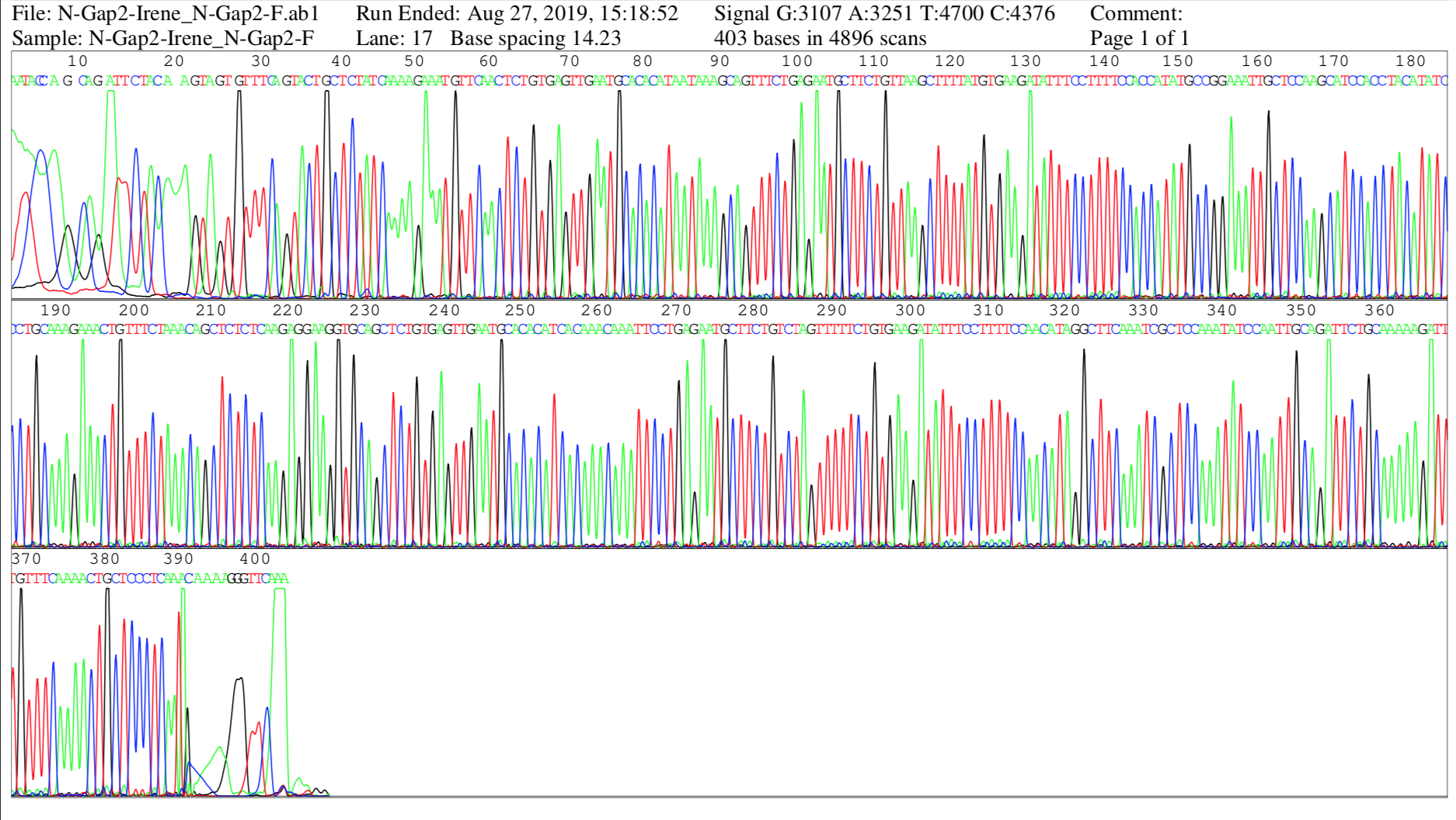

### N-Gap case 2

##### Position (Reference: HG38)

chr20:29,447,139-29,448,514

##### Sequence (N-Gap is marked in red)

ATATTTTGTTCTAAGAATTTTGTAGTTTTAGCTCTTACATTTAGGTATTTGATCCAGTTAGTTAATTTTTTCTTATGGTATAAGTGAATGGCCCAGGTTCATTCTTTTTACATGTGGGTACCGAATTTCCCCAGCCCTGTTCTGCCTCTGACCGTCGGTCCTGTGTGGGTTTACTTGGAGGTGCTTTGCCTTGGAGAAAGGCGGGTGGACGGTGGTGGGGGCAATACAGCCCTGATAACACCTTGACTTAAGCCCAGTAAGACACGTGAGCAACAGGCCCCCTTGCAGAGGGCAAAGCAACGTGGAATCCGAAACCAAGCTTCAGTCGGCCTGAGTGTGACTCCTGTGTGGACGGGACTATCGGCCTCGCGCTCCTTTGCAGGCTCAACCTGGGGCTATCTCATCTGTGAACCATGTGGATGAAAAATGGACAACCACCCGAGTTTCGGCTCATTGCTCTCTGGGCAATTCTCTCATTCCCTGGGAGACGAAATTCGTCTGAATCGCTCCCGGATGAAGCAACCCAGGCTGGTTATCCGGAGGGCCGGTGAGCGCCCCGCAGACCAACGCGGCTGTGGGCCGAGCACTTAGCCCGCACTGGGCACCCAACATTTTTCCGGAGTGCGAGGTCCTGCTGGTCCTGGAGGCAGAAGACCGCTTTTCTCTCTGCCTTCTTTTCTCTGTCCCTTGCTCCCTCCCTCCCTCTTTCCCTCCGTCCCTCCCTCAGTTCCTCCCTTCCTCCCTCCCTCCTTCCCTCTCTTCCTCCCTCTATCCCTCCATCCCTTCCAAGGTCCCTCCGTCCATCCGTTCTTTCCTCCCTCCATCGCTCCCTCCCTCTCTGTCTCCGTTCCTCTCCCCATCTCTGCCTGAGTTCCCTCCCGCGTAGAAAGGGCAGCACCCCGGTTTGCGCGGGGTCTCGGGTCTGCATTTAGCTGTCAGGCGCTCCACGGTGATAGCGAGGAAGCTGGCGGGGCACGGGTAGGCGAGTGACGGTGGCGGGGGGAGGCAGAGGTAGCCAGACGAGGAAAGAAGAGCAGGCCTGGTGGCTGCCCGGGCCAGTGTTTCCCGGGATGGAGGTCTCCGCCCACTCTACTGAAGAACGCGAGTGGGTGGGGGGGAAAGGGATGAGAGCTCCGCCTGCAAACCCGCGCATGAGCAGTAGACGGTTCACCTCCCGGTACCTGGACGGGCTCTGGGATCCCCGGGATGCTCAGGAAAGAATGACAGCCCTCCACTGTGTGGGGTCTCTCACGGGACCTGGAACTCAGGGATCCTAGGCAGGTCAGCTGGAAGGGAAGACACGCCTCTCCATACCGAGTCAGAAGTTCACCGCGAAAGAGAGGCCGCCGCCCCAACCCCGCG

##### Genome Browser

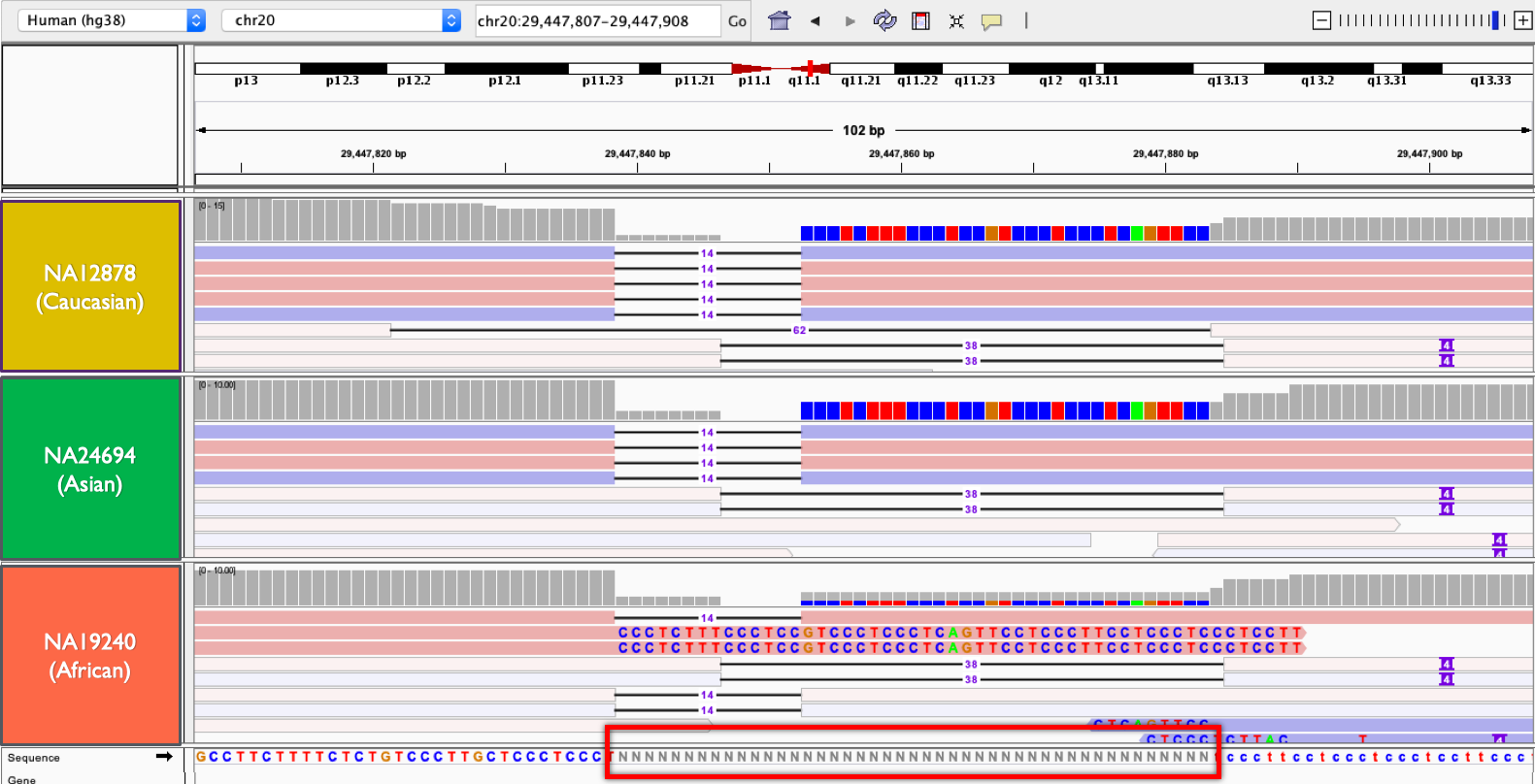

##### PCR Primer

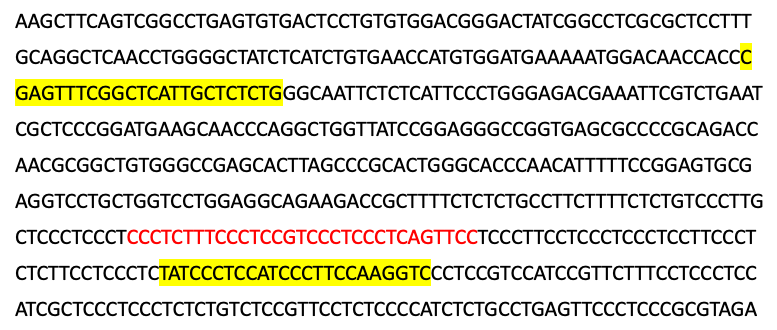

New N-Gap1-F: 5’-CGA GTT TCG GCT CAT TGC TCT CTG-3’(24mer)

N-Gap1-R: 5’-GAC CTT GGA AGG GAT GGA GGG ATA-3’(24 mer)

##### PCR Result

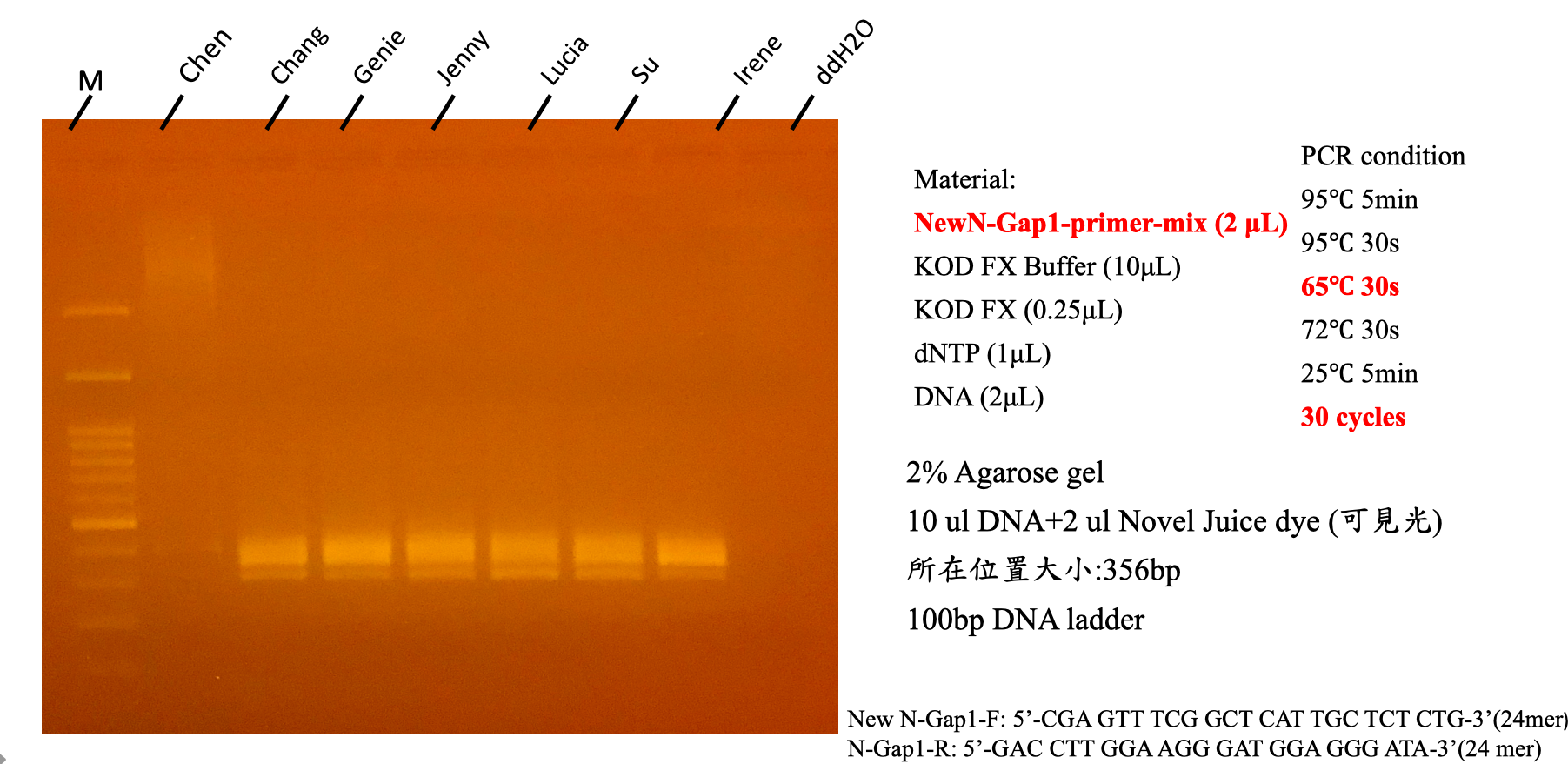

##### Sanger Validation

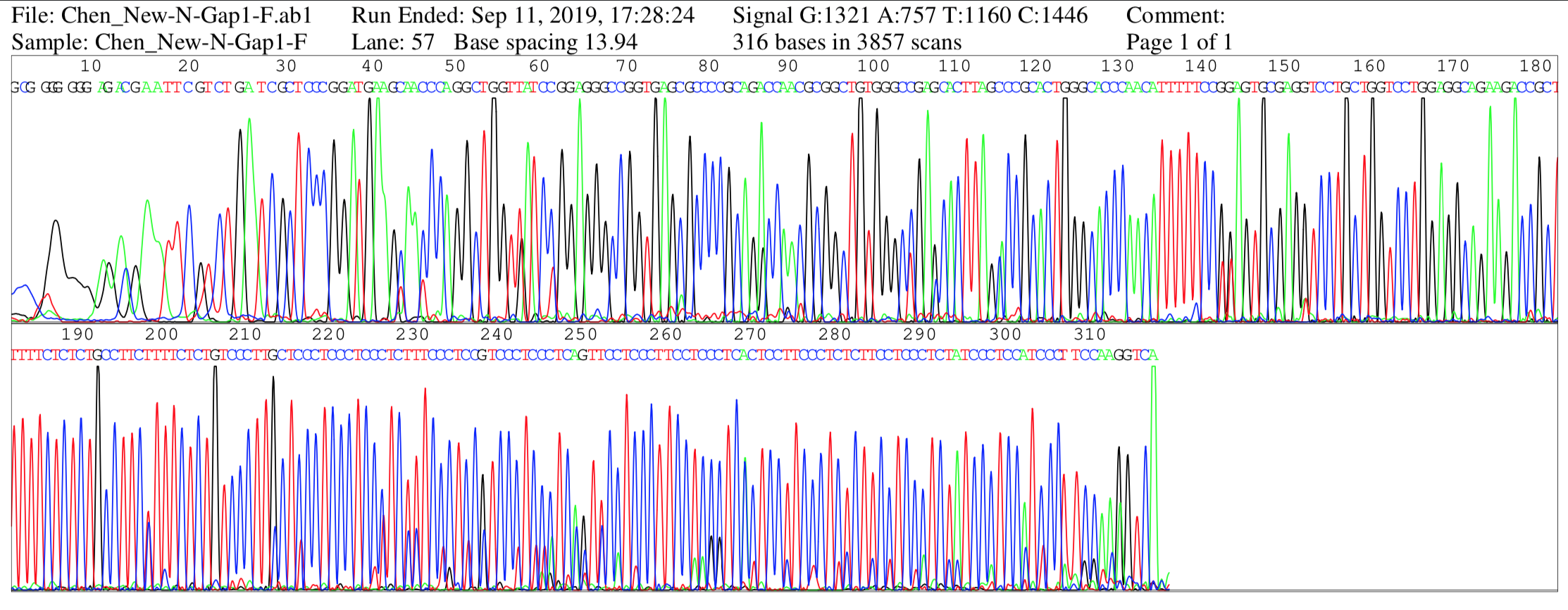

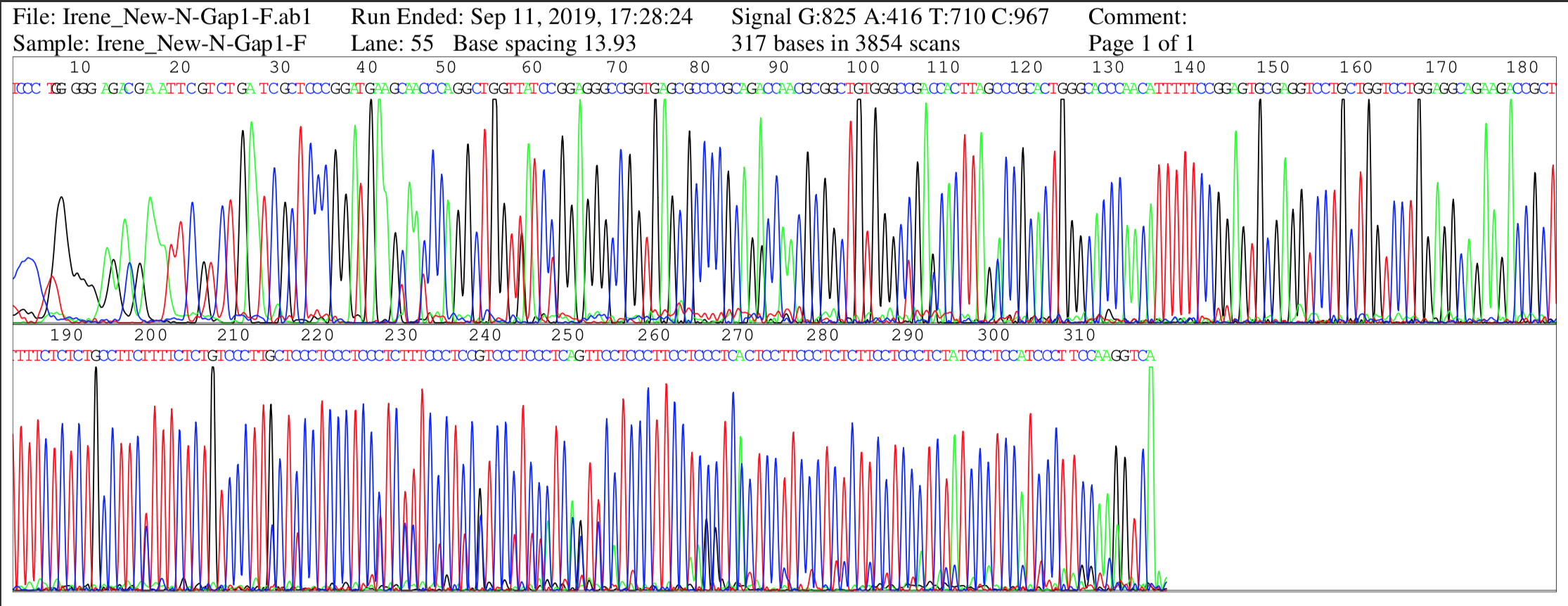
